# *C. elegans* models of Alternating Hemiplegia of Childhood have dominant neuromuscular junction defects

**DOI:** 10.64898/2026.04.22.720250

**Authors:** Diana A. Wall, Adam M. Friedberg, Jeremy Lins, Roza Khalifa, Sienna Partipillo, Anne C. Hart

## Abstract

Dominant missense mutations in ATP1A3, encoding a Na^+^, K^+^ ATPase α-3 subunit, can cause Alternating Hemiplegia of Childhood (AHC), but how these mutations lead to AHC remains unclear. Here, we establish the first *C. elegans* AHC models by introducing AHC-causing ATP1A3 patient mutations (D801N, E815K, L839P, and G947R) into the orthologous gene, *eat-6,* using CRISPR/Cas9. Homozygous *C. elegans* AHC model animals have recessive developmental defects. Heterozygous AHC model animals have dominant defects in neuromuscular junction (NMJ) function that are inconsistent with haploinsufficiency and dominant sleep or arousal defects. Previous work in a *Drosophila* G755S AHC model found that loss of a K⁺-dependent, Na⁺/Ca²⁺ exchanger exacerbated neuronal defects. We introduced a loss-of-function allele of the orthologous *C. elegans* gene, *ncx-4*, into *C. elegans* AHC models; loss of *ncx-4* function did not consistently alter *C. elegans* AHC model defects across alleles. Our results establish novel *C. elegans* models of AHC with robust phenotypes, demonstrate that AHC mutations disrupt NMJ function, and provide proof-of-concept for discovering cross-species modifiers of AHC-related phenotypes.

**Summary Statement:** We report the first *C. elegans* models of Alternating Hemiplegia of Childhood. D801N, E815K, L839P, and G947R AHC model animals have recessive development defects and dominant neuromuscular defects.

## Introduction

Mutations in ATP1A3 cause a spectrum of disorders with overlapping neurological and developmental symptoms (Vezyroglou et al. 2022). Beyond symptomatic treatments, there are no effective therapies for ATP1A3-related diseases, placing a heavy burden on thousands of patients and families (Auvin 2024; Neville and Ninan 2007; Samanta 2020). We focus on dominant, *de novo*, missense mutations in ATP1A3 that cause Alternating Hemiplegia of Childhood (AHC). AHC is classically defined by episodes of paralysis (plegia) that affect one or both sides of the body. Onset occurs in infancy, and patients often present with additional symptoms such as abnormal eye movements, dystonia, seizures, developmental delays, intellectual disabilities, and progressive brain atrophy (Vezyroglou et al. 2022; Heinzen et al. 2012; Rosewich et al. 2012). While over 40 different AHC-associated ATP1A3 mutations have been identified, the most frequently occurring AHC alleles are D801N, E815K, and G947R (Viollet et al. 2015; Heinzen et al. 2012; Verret and Steele 1971; Vezyroglou et al. 2022; Rosewich et al. 2012; Yang et al. 2014; Li et al. 2022). Each of these three AHC alleles is associated with slightly different symptoms. For example, E815K/+ patients are more likely to develop epilepsy, while cardiac muscle function defects are most common in D801N/+ patients (Moya-Mendez et al. 2021; Funk et al. 2025; Moya-Mendez et al. 2025; Panagiotakaki et al. 2015; Viollet et al. 2015; Yang et al. 2014).

ATP1A3 is a catalytic subunit of the P-type Na^+^/K^+^-ATPase that exports Na^+^ and imports K^+^ ions; maintaining appropriate electrochemical Na^+^ and K^+^ gradients is critical for establishing physiologic membrane potential and ionic balance (Kaplan 2002). ATP1A3 is one of four human Na^+^/K^+^-ATPase “alpha” subunits that function as heterodimers with ATP1B family “beta” subunits. Beta subunits are required for maturation, transport to the membrane, and cation pump function of ATP1A proteins (Geering 2001; Kaplan 2002). The impact of AHC patient mutations on ATP1A3 levels, localization, and function has been carefully examined. Patient mutations do not alter ATP1A3 mRNA levels, and most AHC mutations (including D801N and E815K) do not dramatically change ATP1A3 protein levels (Heinzen et al. 2012; Weigand et al. 2014; Blanco-Arias et al. 2009). While AHC alleles do decrease ATP1A3 function, multiple lines of evidence suggest that AHC pathophysiology is more than simple loss of pump activity (Simmons et al. 2018; Sweadner et al. 2019). Other cellular and molecular processes are likely impaired by AHC mutations, leading to dysfunction and patient symptoms.

The *C. elegans* ortholog for all four human ATP1A family proteins is EAT-6 (Davis et al. 1995), which is 72% identical in amino acid sequence to ATP1A3. Complete loss of *eat-6* function is embryonic or early larval lethal, and partial loss-of-function alleles cause recessive defects in pharyngeal pumping, neuromuscular junction (NMJ) function, fecundity, and mechanosensory response (Avery 1993; Davis et al. 1995; Johnson et al. 2020; Doi and Iwasaki 2008; Govorunova et al. 2010; Hawkins et al. 2015). *C. elegans* NMJ function can be readily examined with classical drug-challenge assays using aldicarb, an acetylcholinesterase inhibitor, or levamisole, a nicotinic agonist (J. B. Rand 2007; Lewis, Wu, Levine, et al. 1980; Mahoney et al. 2006; Hobson et al. 2017). Partial loss of *eat-6* function causes recessive hypersensitivity to aldicarb, and some alleles also cause recessive hypersensitivity to levamisole (Govorunova et al. 2010; Doi and Iwasaki 2008). Therefore, decreased EAT-6 function disrupts NMJ function, and some alleles cause post-synaptic defects. EAT-6 function in motor neurons is critical for normal NMJ function (Doi and Iwasaki 2008). For some alleles, EAT-6 function in muscles is critical for normal levamisole response and proper trafficking of post-synaptic nicotinic acetylcholine receptors (Doi and Iwasaki 2008). The deeply conserved sequence and function of mammalian and *C. elegans* ATP1A family proteins support the use of invertebrate models for studying AHC patient mutations.

Here, we report the creation of *C. elegans* AHC models for the most severe and recurrent AHC patient alleles D801N, E815K, and G947R, as well as a less severe AHC allele, L839P. These AHC model animals have recessive developmental defects, dominant arousal defects and dominant NMJ defects. Finally, we use these novel AHC models to test the cross-species relevance of a conserved K^+^-dependent, Na^+^/Ca^+^ exchanger (NCKX), whose loss-of-function modified defects in a *Drosophila* AHC model.

## Results

### Insertion of AHC patient mutations into *C. elegans eat-6*

We selected dominant, missense, ATP1A3 AHC patient mutations D801N, E815K, L839P, and G947R for CRISPR/Cas9-based introduction into endogenous *C. elegans eat-6*. D801N, E815K, and G947R are the most severe and most frequently recurring AHC patient alleles. L839P is a less severe patient allele, which allows for comparison of allele severity between the *C. elegans* models. Each edit to *C. elegans eat-6* was introduced using a single stranded CRISPR repair template designed to include the AHC missense mutation (*e.g.* D801N), as well as silent mutations to remove PAM sequences and create restriction digest sites for PCR genotyping. CRISPR control strains were generated for each AHC allele (*e.g*. D801D) to control for the presence of these silent mutations (Figs. 1, S2, S3). We generated two strains for each AHC allele and for each CRISPR control (See Materials and Methods). Results from the arbitrarily selected “primary” AHC model and CRISPR control strains were very similar to results from the “replicate” AHC model and CRISPR control strains, with minor exceptions described below. Replicate strains are labeled with an asterisk (*). For clarity, we describe the *C. elegans* AHC model strains herein using human amino acid numbering; formal *C. elegans* allele designations can be found in the Supplementary Materials strain list.

**Figure 1.**
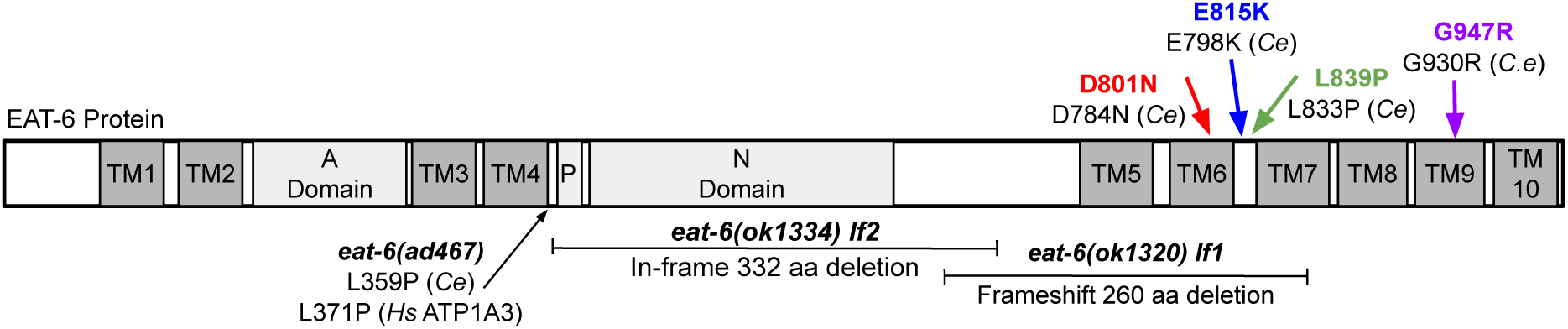
AHC patient mutations and loss-of-function alleles in C. elegans EAT-6. *C. elegans* EAT-6 contains 10 transmembrane (TM) domains, an actuator (A) domain, a phosphorylation (P) site, and a nucleotide binding (N) domain. *eat-6(ad467)* is a previously described homozygous viable, missense allele (Avery, 1993). *eat-6(ok1320)* and *eat-6(ok1334)* are homozygous lethal deletion alleles; in this report *ok1320* is designated *lf1* and *ok1334* is designated *lf2. C. elegans* AHC models for D801N, E815K, L839P, and G947R AHC patient mutations in ATP1A3 are generated in this study by editing endogenous *eat-6*. *Ce* indicates *C. elegans*, and *Hs* indicates *Homo sapiens* amino acid numbering.

### Developmental defects of homozygous AHC model animals

Homozygous AHC model animals had profound developmental defects. These animals either failed to hatch or failed to develop normally (Fig. 2), which was not unexpected as homozygous AHC model animals of other species also have severe homozygous phenotypes (Uchitel et al. 2021; Ng et al. 2021; Hunanyan et al. 2015). Therefore, AHC model animals were maintained as balanced heterozygotes. Heterozygous AHC model animals had one chromosome with the AHC patient mutation in *eat-6* and one balancer chromosome (*tmC12*) with an unedited, functional copy of *eat-6* (*e.g.* D801N/+). Homozygous CRISPR control and heterozygous AHC model animals were healthy and developed into overtly normal, fertile adults without obvious locomotor defects.

**Figure 2.**
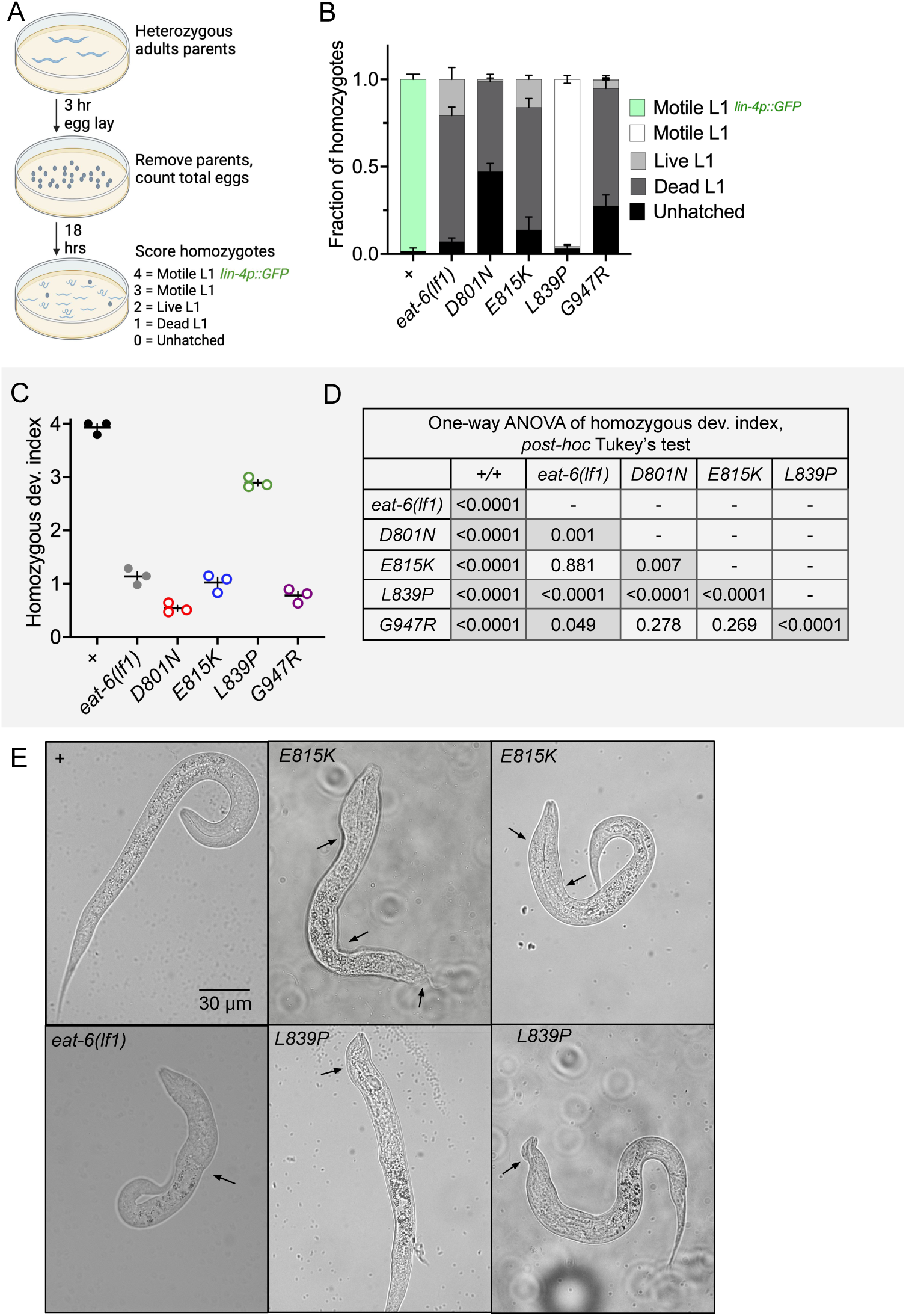
Developmental defects of homozygous AHC model animals. (A) Homozygous development assay and development index used in B and C, respectively. (B) Fraction of homozygous animals 18 hours after egg-lay from balanced, heterozygous parents categorized as unhatched, dead L1, live L1, motile L1, motile L1 expressing *lin-4p::GFP (maIs134)*. n=117-211 homozygous animals per genotype scored from three biological replicates. Error bars indicate mean +/- SEM. (C) Development index score of homozygous animals presented in B. (D) One-way ANOVA of data from C with Tukey’s multiple comparisons test between all pairs. Comparisons in which P<0.05 are shaded. (E) Representative images of homozygous wild-type, E815K, L839P, and *eat-6(lf1)* animals 18 hours after egg-lay. Black arrows point to examples of abnormal body thickness. *eat-6(lf1)* is *ok1320*.

Homozygous AHC model animals are produced by balanced heterozygous parents at the expected Mendelian ratio of 1:3, but homozygous AHC model animals were not viable and never developed past the first (L1) larval stage. These L1 animals were smaller than wild-type, but always had an overtly normal body plan with a pharynx and intestine. Sometimes, homozygous AHC model animals showed abnormal body thinning, bumps, or head swelling (Fig. 2E, black arrows). These phenotypes were highly variable and most frequently observed in homozygous E815K, L839P, and *eat-6* loss-of-function animals.

To carefully examine the developmental defects of homozygous AHC model animals, we recorded the developmental state of homozygous AHC model progeny 18 hours after an egg-lay by heterozygous parents (Fig. 2A-D). Homozygotes were categorized as unhatched, dead L1, live L1, motile L1 that do not reach mid-L1 stage, and motile L1 that do reach mid-L1 stage (Fig. 2A,B). Each category was assigned a developmental index score 0 to 4, respectively, to allow for comparison across lines (Fig. 2A, C-D). To determine if an animal reached the mid-L1 stage, we examined the expression of a previously described developmental marker, *lin-4::GFP (Feinbaum and Ambros 1999; Olsen and Ambros 1999; Ow et al. 2008)*.

Virtually all homozygous wild-type and CRISPR control animals hatched, showed coordinated locomotion, and developed beyond the L1 stage. The *eat-6(ok1320)* loss-of-function allele, referred to as *lf1*, is a 260 amino acid frameshift deletion that removes transmembrane domains 5-7 (Fig. 1). Most homozygous *eat-6(lf1)*, E815K, and G947R animals hatched, but the majority of these L1 larvae were dead. A small fraction of homozygous *eat-6(lf1),* E815K, and G947R larvae were alive: some twitched spontaneously and some only responded when prodded. However, these animals never expressed *lin-4p::GFP* and therefore never developed to the mid-L1 stage. Homozygous D801N model animals had the most severe development defects; almost half of the animals were unhatched and almost all of the L1 larvae were dead (Fig. 2). Homozygous D801N and G947R developmental defects were more severe than homozygous *eat-6(lf1)*, while homozygous E815K defects could not be distinguished from those of homozygous *eat-6(lf1)* animals in this analysis.

Developmental defects observed in homozygous L839P model *C. elegans* were less severe. Almost all of these animals hatched and were motile with slow, spontaneous, sinusoidal movement (Fig. 2). Homozygous L839P model animals lived for more than 10 days, becoming increasingly sessile. We confirmed this for the replicate L839P* homozygous animals (n=30, 100% animals surviving >10 days). By 14 days, about half of the homozygous L839P animals died. Despite their prolonged survival, homozygous L839P animals failed to increase in size and never expressed *lin-4p::GFP* which indicates a failure to develop past the mid-L1 developmental stage. This survival without growth or maturation is similar to that of wild-type animals hatched in the absence of food (Johnson 1983). Combined, these results demonstrate that homozygous *C. elegans* AHC model animals have severe developmental defects, which prevent their growth beyond the earliest stages of development.

### AHC model animals did not have dominant pharyngeal pumping, egg-laying, or stress-response defects

Classical *eat-6* alleles, such as *eat-6(ad467)*, were identified in a forward genetic screen based on recessive pharyngeal pumping defects (Avery 1993). As filter-feeders, *C. elegans* rapidly and rhythmically contract pharyngeal muscles to pump bacteria into the intestine (Avery and Horvitz 1989; Albertson and Thomson 1976), and defective neuromuscular function can decrease pharyngeal pumping rates (Avery 1993; Davis et al. 1995). We examined heterozygous AHC model animals to determine if they also displayed pharyngeal pumping defects. Overall, the *C. elegans* AHC model animals did not consistently show altered pumping rates (Fig. S3A). We note that day-one adults from the D801D*/+ CRISPR control strain had a slightly decreased pumping rate, which means D801N*/+ day one adults appear to have an increased pumping rate in comparison. But this result was not replicated in D801D/+ and D801N/+ animals, whose pumping rates are almost identical (Fig. S3A). We also assessed pharyngeal pumping in day eight adult animals; no consistent pharyngeal pumping defects were observed. We note that E815E*/+ animals had increased pharyngeal pumping rate, when compared to wild-type control animals (Fig. S3B); this was not seen in E815E/+ animals. Both primary and replicate strains of L839L/+ and L839P/+ showed normal pharyngeal pumping rates. Overall, we conclude pharyngeal pumping is not strongly or consistently altered by dominant AHC mutations in *eat-6*.

The standard *E. coli* fed to *C. elegans* is OP50, but rearing *C. elegans* on different *E. coli* strains can reveal subtle feeding defects. DA837 *E. coli* grows in large, sticky clumps and is more difficult for *C. elegans* to ingest (Avery and Shtonda 2003). In contrast, HB101 *E. coli* is low-viscosity and easier for feeding-defective animals to consume (Davis et al. 1995). Heterozygous AHC model animals did not have overt growth rate or fertility changes when grown on DA837 or HB101 *E. coli*. Furthermore, allowing homozygous AHC model progeny to hatch on HB101 *E. coli* did not allow these animals to develop into later stage larvae. Appearance and locomotion of heterozygous AHC model *C. elegans* on all three bacterial strains was indistinguishable from wild-type or CRISPR control strains. We conclude that failure to develop in homozygous AHC model animals was likely not due to feeding defects.

Classical *eat-6* alleles also have recessive egg-laying defects (Davis et al. 1995), as coordinated vulval muscle contractions are required to lay eggs (White et al. 1986). Heterozygous AHC model animals did not show dominant defects in egg-laying (Fig. S3C).

Temperature stress is a common trigger for hemiplegic and dystonic episodes in AHC patients (Heinzen et al. 2014; Salles et al. 2021; Brashear et al., n.d.), and AHC mouse models have temperature-induced paroxysmal episodes (Uchitel et al. 2021; Helseth et al. 2018; Terrey et al. 2025; Isaksen et al. 2017). We exposed heterozygous AHC model *C. elegans* to a variety of acute or chronic heat or cold stress to determine if these could induce dramatic survival or locomotion defects. After a 17 hour 4°C or a 4 hour 0°C cold shock, no survival or locomotion defects were observed in heterozygous D801N/+, E815K/+, or L839P/+ model animals (n=60 animals assessed in 3 biological replicates for each cold shock paradigm, always 100% survival). After a 2.5 hour heat shock at 35°C, heterozygous AHC model animals trended towards a higher survival rate than their CRISPR control strains, but this did not reach significance (Fig. S3D). In combination, these studies suggest that heterozygous AHC model *C. elegans* cannot be distinguished from control animals based on blatant behavioral defects.

### Dominant NMJ defects of AHC model animals: aldicarb hypersensitivity

Animals carrying homozygous viable, partial loss-of-function mutations in *eat-6* have normal locomotion, but drug challenge assays reveal recessive NMJ defects (Doi and Iwasaki 2008; Hawkins et al. 2015; Sorkaç et al. 2016; Johnson et al. 2020; Govorunova et al. 2010). *C. elegans* NMJ function is frequently examined using aldicarb, an acetylcholinesterase inhibitor. In the presence of aldicarb, acetylcholine accumulates at the NMJ and irreversibly induces muscle contraction. Even in wild-type *C. elegans*, exposure to aldicarb will lead to immobilization, due to constant muscle contraction, over time or with increasing dose. In almost every case, an altered rate of immobilization on aldicarb broadly indicates defective NMJ function or developmental defects at the NMJ. Note: aldicarb-induced immobilization is not directly equivalent to dystonia or paralysis observed in AHC patients but does serve as a robust assessment of overall NMJ function. The aldicarb assay alone cannot determine if NMJ dysfunction results from defects in neurotransmitter (acetylcholine and/or GABA) release and/or the post-synaptic response by muscles. (James B. Rand 2007; Mahoney et al. 2006). We used two different aldicarb assay formats to examine NMJ function in heterozygous *C. elegans* AHC model animals: dose-response and time-response. We found that *C. elegans* AHC model animals had dominant NMJ defects in both assay formats.

In the dose-response aldicarb assay, we measured the fraction of animals immobilized after five hours on 0, 0.25, 0.50, 0.75, and 1 mM aldicarb (Fig. 3A). In this assay, heterozygous *eat-6(lf1)/+* animals responded normally to aldicarb (based on both two-way ANOVA versus balanced wild-type animals, and on area under the curve (AUC) values using a paired t-test, Fig. 3B,C). Primary and replicate CRISPR control strains also responded normally to aldicarb when compared to balanced, wild-type animals (Fig. 3D-F). All eight primary and replicate heterozygous AHC model strains showed dominant hypersensitivity to aldicarb when compared to their heterozygous CRISPR control strains (Fig. 3D-R). The aldicarb hypersensitivity of all AHC model strains was seen in both two-way ANOVA of the dose-response curves and in paired t-tests of AUC values. This indicates all four *C. elegans* AHC models show dominant defects in NMJ function, while *eat-6(lf1)/+* animals do not.

**Figure 3.**
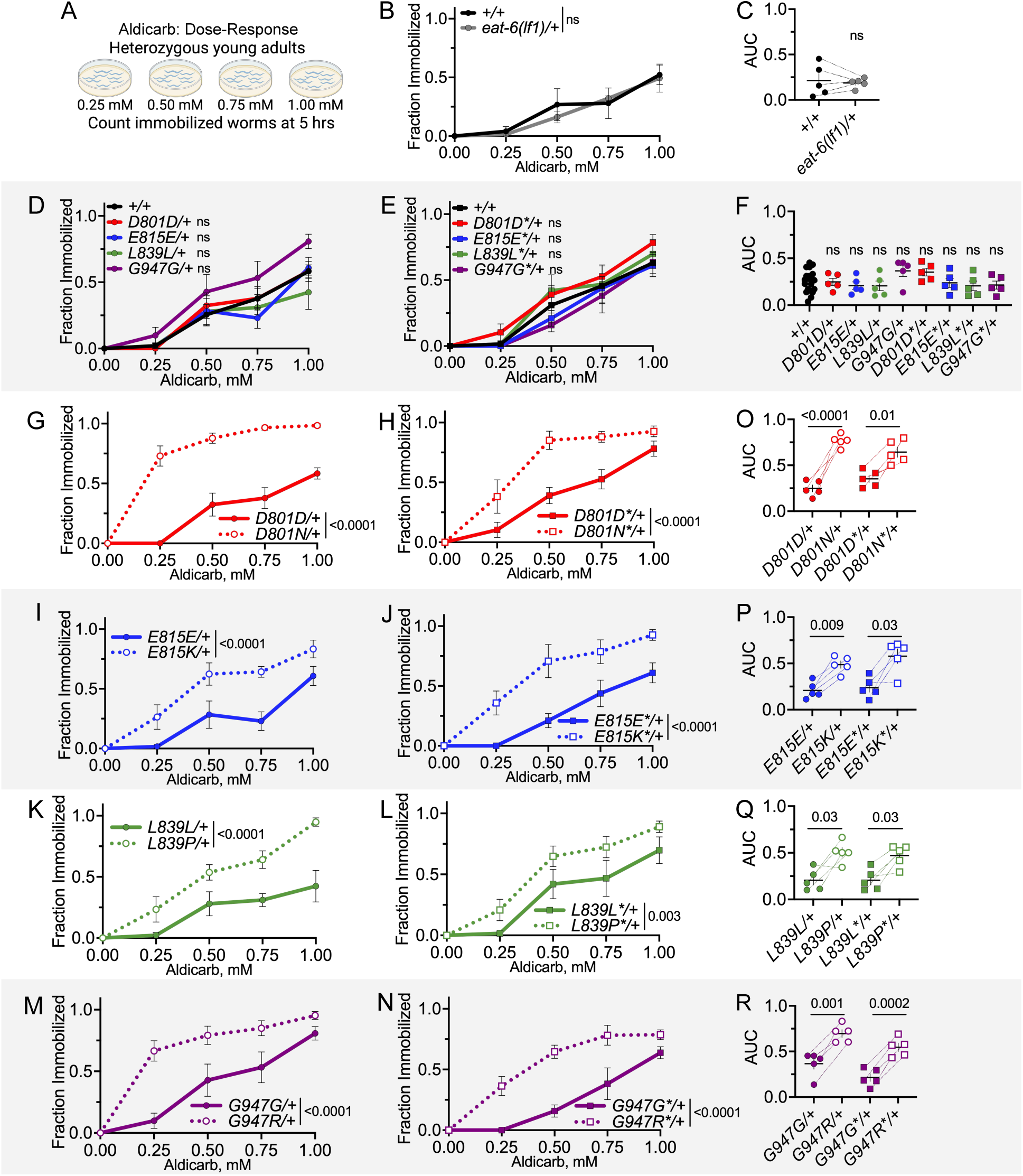
Dominant NMJ defects of AHC model animals: aldicarb dose-response. (A) Aldicarb dose-response assay. The number of heterozygous, young-adult animals immobilized on 0, 0.25, 0.5, 0.75, and 1 mM aldicarb was recorded at five hours. (B) Dose response curve for *eat-6(lf1)/+* animals; *lf1* is *ok1320*. Two-way ANOVA. (C) Area under the curve (AUC) from each trial in B. Paired t-test. (D,E) Dose response curves for heterozygous CRISPR control primary and replicate strains. Accumulated +/+ data is plotted for visual purposes, but two-way ANOVA was tested for CRISPR control and +/+ animals run in parallel. (F) AUC from each trial in D and E. Paired t-test between each CRISPR control and +/+ animals run in parallel. (G-N) Dose response curves for heterozygous primary and replicate CRISPR control and AHC model strains. CRISPR control data is the same as in D,E. Two-way ANOVA between each CRISPR control and AHC model strain. (O-P) AUC for each trial in G-N. Paired t-test between CRISPR control and AHC model strains tested in parallel. For all two-way ANOVA analysis, the displayed P value results from genotype as the source of variation. ns indicates P>0.05. All animals in B-R have one copy of the *tmC12* balancer represented by “+”. Each genotype tested in five biological replicates. For each trial, n=10-15 animals per genotype per dose. Error bars indicate mean +/- SEM.

Different assay formats can reveal subtle differences between genotypes. Therefore, we also assessed NMJ function in an aldicarb time-response format. Here, we measured the cumulative fraction of animals immobilized on 1mM aldicarb hourly over the course of six hours (Fig. 4A). Again, we found that heterozygous loss of *eat-6* function did not cause dominant aldicarb hypersensitivity (Figs. 4B, 3C, *eat-6(ok1320 lf1)/+* and *eat-6(ok1334 lf2)/+* animals, based on Mantel-Cox log-rank test and paired t-test of median immobilization time). Heterozygous primary CRISPR control strains were not different from wild-type animals (Fig. 4D). We note that replicate D801D*/+ and L839L*/+ animals were hypersensitive to aldicarb in this assay format when compared to balanced wild-type animals based on a Mantel-Cox log-rank test (Fig. 4E). When median immobilization time is examined, no CRISPR control primary or replicate strain differs from wild-type. Background mutations in D801D*/+ and L839L*/+ strains likely have a small effect on NMJ function.

**Figure 4.**
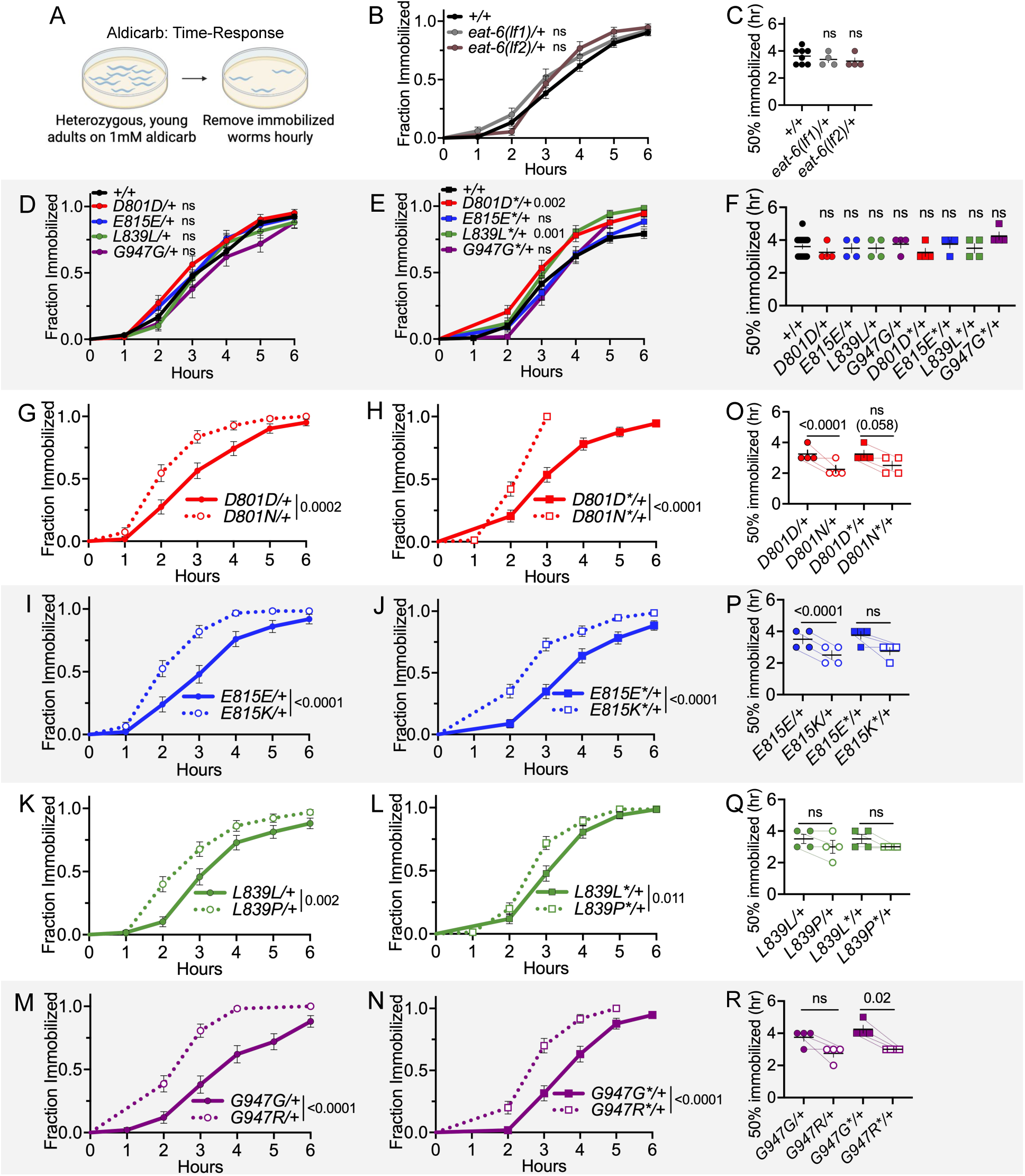
Dominant NMJ defects of AHC model animals: aldicarb time-response. (A) Aldicarb time-response assay. The number of heterozygous, young-adult animals immobilized on 1 mM aldicarb was recorded hourly. (B) Immobilization curve for *eat-6(ok1320)/tmC12* (*lf1*/+) and *eat-6(ok1334)/nT1* (*lf2*/+) and balanced wild-type animals. For visual purposes, +/+ represents aggregated immobilization curves of both *+/nT1* and *+/tmC12* animals. Mantel-Cox log-rank analysis performed between each balanced *eat-6* allele and corresponding wild-type balanced strain tested in parallel. (C) Median immobilization time for each trial in B. Paired t-test between each balanced *eat-6* allele and corresponding wild-type control. (D,E) Immobilization curves for heterozygous CRISPR control primary and replicate strains. +/+ data is aggregated for visual purposes, but Mantel-Cox log-rank analysis performed on CRISPR control and +/+ animals tested in parallel. (F) Median immobilization time for each trial in D and E. Paired t-test between CRISPR control and +/+ strains run in parallel. (G-N) Immobilization curve for heterozygous primary and replicate CRISPR control and AHC model strains. CRISPR control data is the same as in D,E. Mantel-Cox log-rank analysis between CRISPR control and AHC model strain tested in parallel. (O-P) Median immobilization time for each trial in G-N. Paired t-test between CRISPR control and AHC model strains run in parallel. Each genotype tested in four biological replicates. For each trial, n=10-22 animals per genotype. Error bars indicate mean +/- SEM. ns indicates P>0.05. All strains in D-R have one copy of the *tmC12* balancer represented by “+”.

In the time-response assay format, heterozygous animals from all eight primary and replicate AHC model strains still showed a dominant hypersensitivity to aldicarb; they immobilized faster than their CRISPR control strains (based on Mantel-Cox log-rank test, Fig. 4G-N). When assessing these results using paired t-tests of median immobilization time, animals from only one of the two independent strains showed dominant aldicarb hypersensitivity for D801N/+, E815K/+, and G947R/+ while neither of the L839P/+ animals showed dominant defects. The Mantel-Cox log-rank analysis may more robustly detect aldicarb hypersensitivity as it takes into account the entire immobilization curve, whereas the median immobilization rate is a single parameter. We conclude that inserting AHC patient missense mutations into *eat-6* causes dominant NMJ defects, based on changes to aldicarb sensitivity in heterozygous AHC model animals in two different assay formats. We also conclude that AHC patient mutations do not cause dominant NMJ defects in *C. elegans* due to haploinsufficiency since heterozygous *eat-6(lf)/+* animals always responded normally to aldicarb.

The magnitude of aldicarb hypersensitivity in heterozygous AHC model animals is similar to that of homozygous *eat-6(ad467)* partial loss-of-function animals (Fig. S6). However, since aldicarb hypersensitivity is recessive in homozygous *eat-6* partial loss-of-function animals and dominant in heterozygous AHC model animals, we do not gain further insight regarding the nature of AHC mutations solely based on this comparison. The *eat-6* partial loss-of-function mutation and the AHC patient mutations could cause aldicarb hypersensitivity through very different mechanisms.

### Dominant post-synaptic NMJ defects of AHC model animals: levamisole hypersensitivity

Levamisole, an acetylcholine receptor agonist, can reveal post-synaptic NMJ defects in *C. elegans*. Similarly to aldicarb, exposure to levamisole causes immobilization of *C. elegans.* Levamisole mimics acetylcholine to constitutively activate receptors on post-synaptic muscles. In mutant animals, an abnormal rate of immobilization on levamisole indicates defective post-synaptic body wall muscle function (Lewis, Wu, Berg, et al. 1980; Lewis, Wu, Levine, et al. 1980). Here, post-synaptic body wall muscle function was assessed using a levamisole time-response assay in which the cumulative fraction of heterozygous animals immobilized on 100 µM levamisole hourly over the course of six hours (Fig. 5A). Heterozygous *eat-6(lf1)/+* animals responded normally to levamisole (based on Mantel-Cox log-rank test and paired t-test of median immobilization time, Fig. 5B,C). When compared to balanced wild-type controls, D801D/+, L839L/+, and all four replicate CRISPR control strains were resistant to levamisole based on a Mantel-Cox log-rank test (Fig. 5D,E). Only G947G*/+ control animals differed from wild-type, based on a paired t-test of median immobilization times (Fig. 5F). These defects may be due to background mutations and confirms the importance of comparing AHC model animals to their respective CRISPR control strain, especially in a levamisole-sensitivity assay. Heterozygous animals from both primary and replicate heterozygous AHC model strains for D801N/+, L839P/+, and G947R/+ showed hypersensitivity to levamisole, when compared to their CRISPR controls, based on a Mantel-Cox log-rank test (Fig. 5G-N). The replicate heterozygous E815K*/+ strain was also hypersensitive to levamisole when tested this way; only heterozygous E815K/+ animals responded normally to levamisole in the log-rank test (Fig. 5I,J). When median immobilization times were compared in paired t-tests, only D801N/+, D801N*/+, and G947R*/+ model animals showed levamisole response defects (Fig. 5O-R). This may indicate that the levamisole response defects in *C. elegans* D801N/+, and perhaps G947R/+, AHC model animals are more robust.

**Figure 5.**
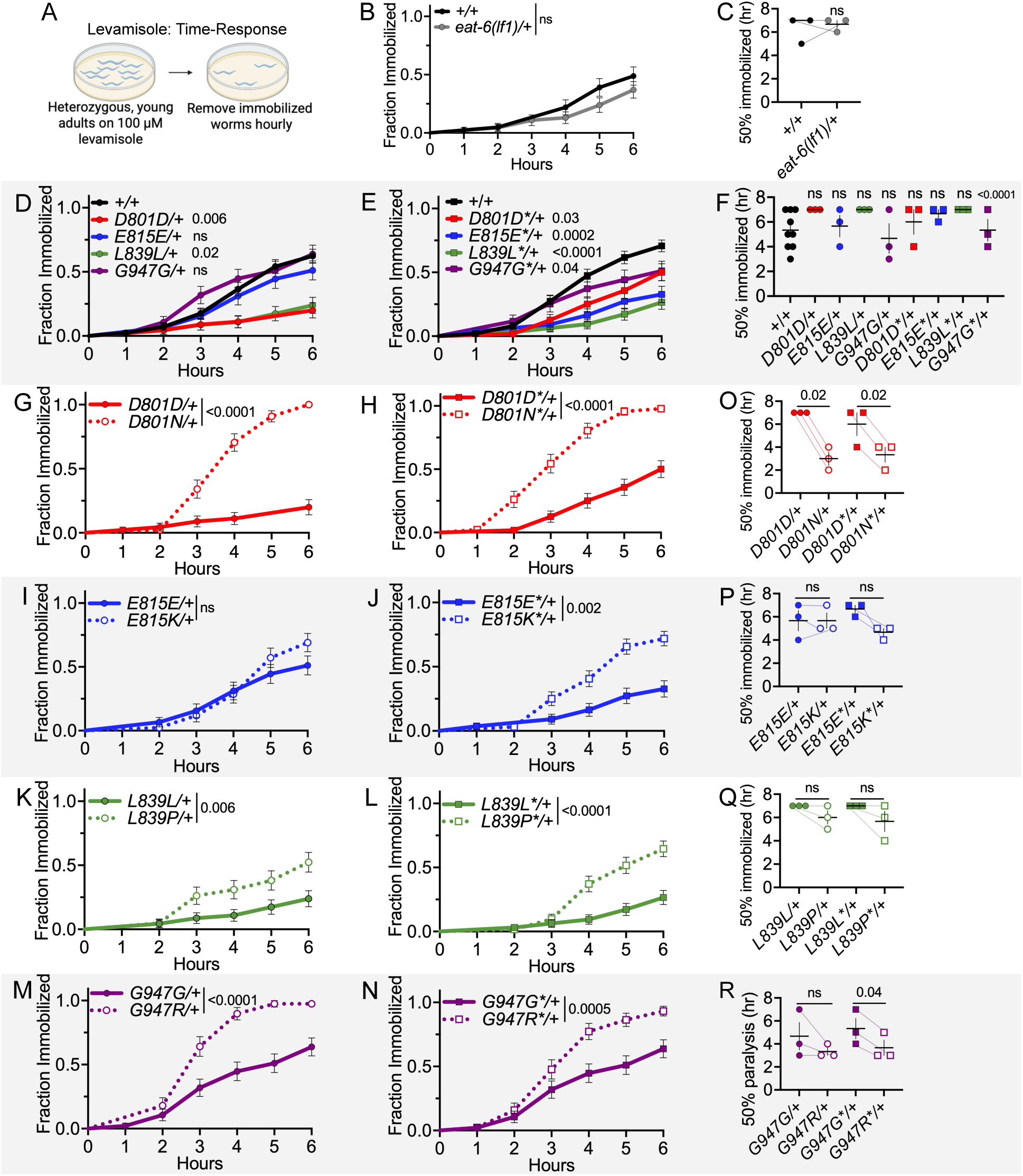
Dominant post-synaptic NMJ defects of AHC model animals: levamisole time-response. (A) Levamisole time-response assay. The number of heterozygous, young adult animals immobilized on 100 uM levamisole was recorded hourly. (B) Immobilization curve for heterozygous *eat-6(lf1), ok1320,* and +/+ animals. Mantel-Cox log-rank analysis. (C) Median immobilization time for each trial in B. Paired t-test. (D,E) Immobilization curves for +/+ and heterozygous CRISPR control primary and replicate strains. +/+ data is aggregated for visual purposes, but Mantel-Cox log-rank analysis performed on CRISPR control and +/+ animals tested in parallel. (F) Median immobilization time for each trial in D, E. Paired t-test between CRISPR control and +/+ strains tested in parallel. (G-N) Immobilization curve for heterozygous primary and replicate CRISPR control and AHC model strains. CRISPR control data is the same as in D,E. Mantel-Cox log-rank analysis between CRISPR control and AHC model strains tested in parallel. (O-P) Median immobilization time for each trial in G-N. Paired t-test between CRISPR control and AHC model strains tested in parallel. Data for each genotype is from three biological replicates. For each trial, n=9-27 animals per genotype. Error bars indicate mean +/- SEM. ns indicates P>0.05. All strains in D-R carry the *tmC12* balancer represented by “+”.

Overall, we conclude that all *C. elegans* AHC models show dominant NMJ function defects. For all strains except the primary E815K/+, this includes post-synaptic body-wall muscle defects. It is important to note that, while levamisole hypersensitivity demonstrates that a post-synaptic muscle defect is present, this does not rule out simultaneous defects in the pre-synaptic neurons that can also contribute to the NMJ defects. Furthermore, we conclude that the dominant post-synaptic muscle defects in heterozygous AHC model animals are caused by a mechanism other than haploinsufficiency, as heterozygous *eat-6(lf1)/+* animals respond normally to levamisole.

### Dominant sleep and arousal defects of *C. elegans* AHC model animals

Many AHC patients experience irregular sleeping patterns (Ricci 1995; Kansagra et al. 2019). Many of the molecular pathways that regulate sleep and arousal are conserved across the animal kingdom, and *C. elegans* exhibit all of the hallmark behavioral changes of sleep during developmentally-timed sleep (DTS), which occurs after every larval stage (Trojanowski and Raizen 2016; Singh et al. 2014). During DTS, animals enter short motionless “sleep bouts” interspersed with “waking bouts” over the course of several hours, in a period called lethargus. Here, we examine sleep in heterozygous AHC model animals during the final lethargus period, as animals transition between L4 and adulthood (L4/adult lethargus). We found that heterozygous G947R/+ animals spent less total time asleep (Fig. 6A), which can be explained by their shorter lethargus duration (Fig. 6B), compared to G947G/+ control animals. Sleep quantity and lethargus duration were normal in heterozygous D801N/+ and E815K/+ model animals, compared to control strains (Fig. 6B,C). However, E815K/+ animals entered lethargus 2.2 +/- 0.8 hours later than E815E/+ control animals (95% CI [0.64, 3.7], Fig. 6D), which may indicate developmental delay. There was no change in the average sleep bout duration for any of the heterozygous AHC model strains tested (Fig. S4). We conclude that heterozygous G947R/+ animals have dominant defects in sleep quantity and heterozygous E815K/+ animals have delayed sleep onset.

**Figure 6.**
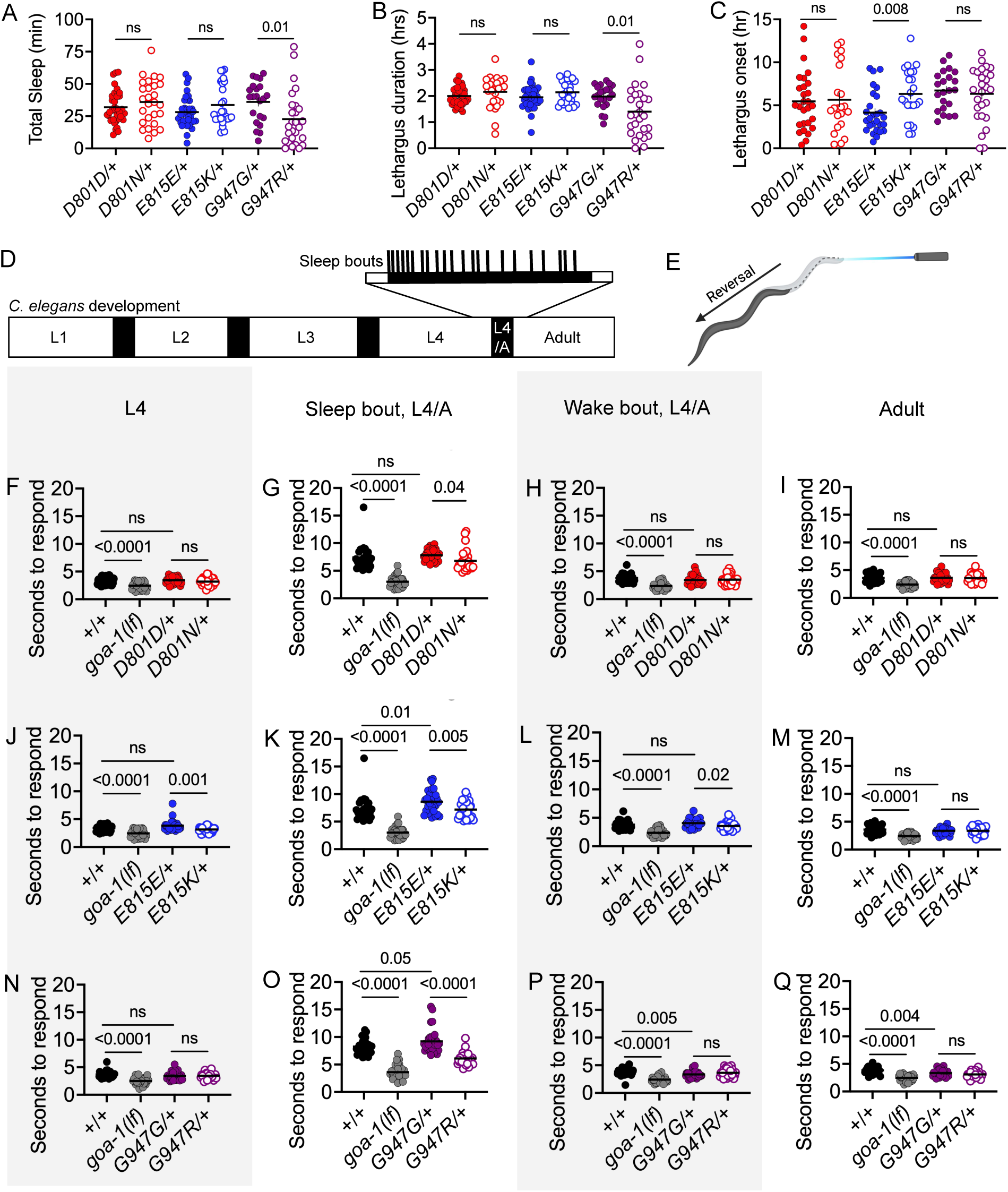
Dominant sleep and arousal defects in C. elegans AHC model animals. (A) Total sleep (minutes), (B) lethargus duration (hours), (C) lethargus onset time (hour) recorded in heterozygous CRISPR control and AHC model animals during developmentally-timed sleep (DTS) during lethargus at the L4 larval to adult stage transition (L4/A). Welch’s unpaired t-test between each CRISPR control and AHC model strain. Each AHC model strain was assessed in separate trials, but presented together here for visual purposes. n= 21-34 animals per genotype. (D) *C. elegans* larval and adult stages (white boxes), separated by lethargus which occurs between stages (black boxes). Results in A-C are from L4/A lethargus. Arousal data in F-Q was collected at the L4, L4/A lethargus, or adult stage. (E) *C. elegans* arousal assay. The time to reverse in response to blue light is recorded. (F-Q) Time to respond to blue light for L4, sleep bout during L4/A, wake bout during L4/A, and adult animals. One-way ANOVA with Sidak’s multiple comparisons test between pairs. *goa-1(sa734lf)* is an internal control for decreased arousal response. n=30 animals per genotype, tested in three biological replicates. For D801D/+, D801N/+, E815E/+, and E815K/+ animals, only “+” chromosome is the *nT1* balancer. For G947G/+ and G947R/+ animals, “+” chromosome represents the *tmC12* balancer. +/+ animals are the N2 wild type strain. Error bars indicate mean +/- SEM.

During sleep, arousal thresholds increase which can be observed by a slower response to mild stimulation, compared to awake animals. We measured arousal thresholds in heterozygous AHC model animals at four times: L4 larval stage, sleep bout during L4/adult lethargus, wake bout during L4/adult lethargus, and adult. Arousal thresholds were determined by measuring the time required for a response to an aversive blue-light stimulus (Edwards et al. 2008; Huang et al. 2018). Shorter response times before and after lethargus suggest a global hyperarousal state; whereas shorter response times only during lethargus sleep bouts suggest poor sleep quality. We found that heterozygous D801N/+, E815K/+, and G947R/+ model animals were all easier to rouse during sleep bouts in L4/adult lethargus, compared to their CRISPR controls (Fig. 6G,K,O), suggesting poor sleep quality. At the adult stage, response times for all AHC models did not differ from their controls, suggesting that AHC model animals are not in a global hyperarousal state (Fig. 6I,M,O). In addition to their lethargus sleep bout defects, heterozygous E815K/+ animals also showed arousal defects during L4 larval stage and L4 lethargus wake bouts (Fig. 6J,L). This, along with the delayed lethargus onset observed in E815K/+ animals, suggests that these animals may have more difficulty transitioning from a wake-state to a sleep-state. The dominant arousal defects reported here indicate that sleep is not normal in heterozygous *C. elegans* AHC model animals, even when the quantity of sleep (defined by lack of motion) does not change during lethargus. Each AHC model shows unique defects when sleep and arousal are examined.

### NMJ defects in *C. elegans* AHC model animals lacking NCX-4 function

ATP1A family protein function is deeply conserved across species; we consider it likely that cellular mechanisms affected in AHC-related defects are similarly conserved. To date, no human genetic modifiers of AHC severity, onset, or outcome have been identified, so we turned to modifier genes identified in another animal model. The *Atpα* gene encodes the *Drosophila* ortholog of all four human ATP1A subunits and *C. elegans* EAT-6. In 2014, the Palladino group reported results from a genome-wide *Drosophila* screen that identified genes whose decreased function modified behavioral defects of three different dominant *Atpα* mutations (Talsma et al. 2014). Providentially, one of the dominant *Atpα* alleles used in the *Drosophila* screen, *CJ10*, is a *Drosophila* model for the AHC patient allele G755S, which was first reported in the literature that same year (Ashmore et al. 2009; Sasaki et al. 2014). The Palladino group identified 33 modifier genes whose decreased function altered the bang-senstive paralysis defects of heterozygous G755S *Drosophila* AHC model animals (Talsma et al. 2014). Determining which *Drosophila* modifier genes encode cross-species modifiers of ATP1A family proteins should reveal conserved pathways directly relevant to AHC dysfunction. Here, we report cross-species analysis of one *Drosophila* modifier, *Nckx30C,* whose decreased function exacerbated *Drosophila CJ10* dominant bang-sensitive paralysis*. Nckx30C* encodes a transmembrane K^+^-dependent Ca^2+^/Na^+^ exchanger (NCKX) that is primarily expressed in adult *Drosophila* neurons (Winkfein et al. 2004). Based on amino acid similarity, the likely *C. elegans* ortholog of *Nckx30C* is *ncx-4*, and the likely human ortholog is *SLC24A2*. We obtained two loss-of-function deletion alleles for *ncx-4*; *ncx-4(tm5106)* and *ncx-4(tm5296)* referred to here as *ncx-4(lf1)* and *ncx-4(lf2)*, respectively (Fig. 7A).

**Figure 7.**
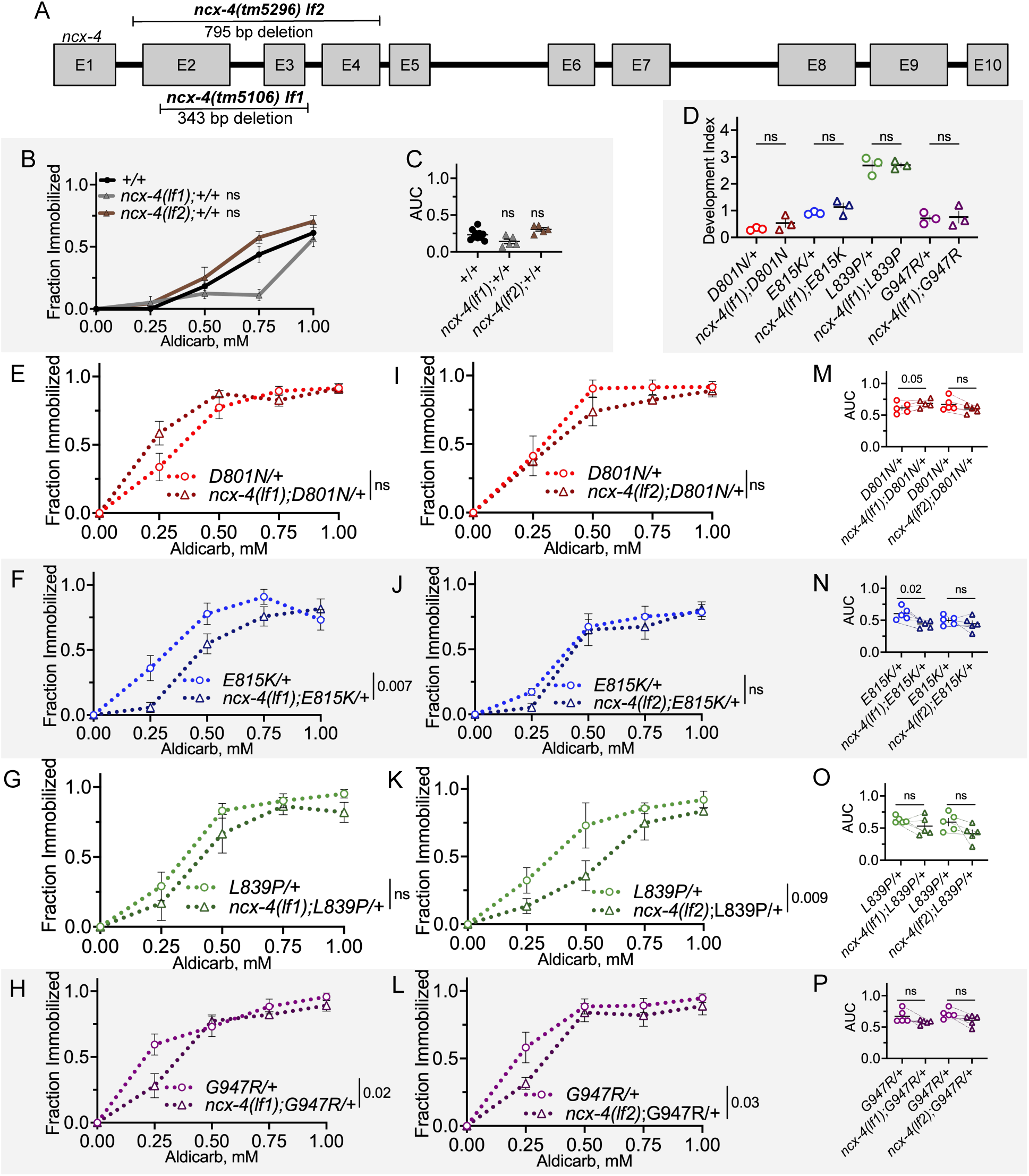
NMJ defects in C. elegans AHC model animals lacking ncx-4 function. (A) *ncx-4(tm5106)* and *ncx-4(tm5296)* are loss of function deletion alleles, designated here as *lf1* and *lf2.* (B) Aldicarb dose-response curve for *ncx-4*(*lf1);+/+ and ncx-4(lf2);+/+* animals. Each *ncx-4* allele was tested separately, and +/+ data from both experiments are aggregated for presentation. Two-way ANOVA analysis performed on *ncx-4* loss of function and +/+ animals tested in parallel. (C) Area under the curve (AUC) values for each trial in B. Paired t-test between animals tested in parallel. (D) Development index of homozygous AHC model animals with and without *ncx-4(lf1)*. Welch’s unpaired t-tests between animals with and without *ncx-4(lf1)* of the same AHC allele. Data collected from three biological replicates. n=114-206 homozygotes per genotype scored. (E-H) Aldicarb dose-response curves for heterozygous AHC model animals with and without homozygous *ncx-4(lf1)*. Two-way ANOVA between genotypes tested together. (I-L) Aldicarb dose-response curves for heterozygous AHC model animals with and without homozygous *ncx-4(lf2)*. Two-way ANOVA between genotypes tested together. (M-P) AUC values from each trial in E-L. Paired t-tests between genotypes tested together. For all two-way ANOVA analysis, displayed P value results from genotype as the source of variation. ns indicates P>0.05. In all animals, one wild-type “+” chromosome represents the *tmC12* balancer. Data for each genotype collected from five biological replicates. For each trial, n=13-15 animals per genotype. Error bars indicate mean +/- SEM.

We first determined whether loss of *ncx-4* function altered the developmental defects in homozygous *C. elegans* AHC model animals. Homozygous AHC model animals also homozygous for *ncx-4(lf1)* showed no alteration to developmental defects; growth defects were not ameliorated or exacerbated (Fig. 7D).

We found that in heterozygous *C. elegans* AHC model animals, heterozygous loss of *ncx-4* did not alter the dominant aldicarb hypersensitivity of heterozygous D801N/+ or E815K/+ animals (*ncx-4(lf1)/+,* Fig. S5D-G). For the remainder of the *ncx-4* studies, the impact of homozygous loss of *ncx-4* function was examined. Homozygous loss of *ncx-4* function alone did not alter aldicarb sensitivity, compared to wild-type control animals (two-way ANOVA, Fig. 7B and paired t-test of AUC, Fig. 7C).

When we introduced homozygous *ncx-4* loss-of-function alleles into heterozygous AHC models, the effect size was not large and results varied by allele. Complete loss of *ncx-4* did not affect aldicarb hypersensitivity of D801N/+ animals based on a two-way ANOVA (Fig. 7E,I), but an exacerbation of NMJ defects was detected only in *ncx-4(lf1);D801N/+* animals based on paired t-test of AUC values (Fig. 7M). Aldicarb hypersensitivity was suppressed by complete loss of *ncx-4* in G947R/+ animals; this suppression was observed for both *ncx-4(lf1)* and *ncx-4(lf2)* loss-of-function alleles (two-way ANOVA, Fig. 7H,L). However, when AUC values were assessed in paired t-tests, neither *ncx-4(lf1)* nor *ncx-4(lf2)* modified G947R/+ animal aldicarb hypersensitivity. In E815K/+ animals, *ncx-4(lf1)* suppressed NMJ defects (two-way ANOVA, Fig. 7F, and AUC paired t-test, Fig. 7N), while *ncx-4(lf2)* had no impact. Finally, in L839P/+ animals, only *ncx-4(lf2)* suppressed NMJ defects (two-ANOVA, Fig. 7K, but not AUC pair t-test, Fig. 7O), while *ncx-4(lf1)* had no effect. Overall, we note that the effect size of *ncx-4* was not large in any case; if *ncx-4* is a cross-species modifier, then it is a weak one. The consistent suppression of NMJ defects in G947R/+ animals when *ncx-4* function is lost could reflect unique defects caused by this AHC allele, but the magnitude of the effect is not large. Nevertheless, these results demonstrate that dominant NMJ defects in *C. elegans* AHC model animals can be enhanced or suppressed, and there may be conserved genetic mechanisms relevant to AHC defects.

## Discussion

### Development of the first *C. elegans* models of Alternating Hemiplegia of Childhood

We created the first *C. elegans* models of Alternating Hemiplegia of Childhood through direct editing of endogenous *eat-6*, which encodes the *C. elegans* ortholog of human ATP1A family proteins. Our models include the three most frequently recurring AHC-associated ATP1A3 mutations in AHC patients (D801N, E815K, G947R) as well as a rare and milder variant (L839P), allowing for comparison of allele-specific phenotypes. Our findings demonstrate that introducing AHC patient missense mutations into *eat-6* causes recessive developmental defects and dominant NMJ defects. Based on abnormal responses to levamisole, post-synaptic muscle defects may contribute to NMJ defects. The developmental, behavioral and NMJ defects reported herein establish *C. elegans* models of AHC, which can be used as patient genotype archetypes to investigate conserved disease mechanisms, as well as to inform the allele-specific properties and genotype-phenotype relationships of disease-associated ATP1A3 variations.

Current model systems for AHC include HEK293 cell lines, *Xenopus* oocytes, mouse models, and patient-derived iPSC neurons (Arystarkhova et al. 2019, 2021; Ng et al. 2021; Uchitel et al. 2021; Li et al. 2015; Simmons et al. 2018). Unfortunately, none of the current *in vitro* assays correlate with allele-specific disease severity (Lazarov et al. 2020), and generating mouse models for the numerous AHC-associated mutations in ATP1A3 would be laborious and expensive. The *C. elegans* AHC models are complementary to the current AHC model systems and can immediately be used for compound screening, identification of genetic suppressors, or testing hypotheses regarding allele-specific suitability of different therapeutic approaches.

When these studies are undertaken, best practice will be to compare results in heterozygous *C. elegans* AHC models to their heterozygous CRISPR controls, rather than using balanced wild-type animals as controls. This is especially important in levamisole-sensitivity assays as multiple CRISPR control strains were resistant to levamisole when compared to wild-type (Figs. 5D,E). As using CRISPR control strains may not be convenient in all assay formats (e.g. large-scale screening), we re-analyzed the aldicarb and levamisole sensitivity data from Figures 3, 4 and 5 comparing balanced heterozygous AHC animals to balanced wild-type “+/+” animals that were tested in parallel (Supplementary Materials Raw Data file). For aldicarb assays, interpretations do not change; all primary and replicate heterozygous AHC model strains still show robust dominant aldicarb hypersensitivity in a two-way ANOVA or log-rank test. For levamisole sensitivity assays, the post-synaptic defects observed in E815K/+ and L839P/+ animals are no longer significant, when compared to balanced wild-type “+/+” animals as a control in a log-rank test. Combined, these results suggest that background mutations may have been introduced during CRISPR editing, or that the “silent edits” in CRISPR control strains could impact phenotypes in some assays, as we sometimes detected differences between balanced CRISPR control and balanced wild-type animals (Figs. 4E, 5D,E, 6K,O,P,Q).

### Dominant NMJ defects in *C. elegans* AHC models are inconsistent with haploinsufficiency

It is well established that dominant AHC mutations in ATP1A3 decrease Na^+^/K^+^-ATPase activity (Li et al. 2015; Simmons et al. 2018; Immanneni et al. 2024). However, multiple lines of evidence suggest that AHC mutations cause additional antagonistic effects which contribute to AHC pathophysiology. D801N, E815K, and G947R mutations in ATP1A3 have dominant-negative effects on wild-type ATP1A3 activity in *Xenopus* oocytes (Li et al. 2015). In HEK293 cell lines, expression of AHC-associated ATP1A3 variants cause immature ATP1B1 “beta” subunits to accumulate in the endoplasmic reticulum and/or Golgi (Sweadner et al. 2019; Arystarkhova and Sweadner 2024; Arystarkhova et al. 2021). In *C. elegans*, EAT-6 also plays a role in the expression and localization of post-synaptic receptors in a manner that is independent from ATPase activity (Doi and Iwasaki 2008), which could help explain NMJ defects observed in AHC model animals. Lastly, the behavioral defects in mice heterozygous for ATP1A3 knockout are not as severe as the defects of heterozygous AHC model mice, suggesting a mechanism beyond simple loss of ATPase function contributes to defects (Sugimoto et al. 2018; Terrey et al. 2024).

The *C. elegans* AHC models described here provide additional support for the hypothesis that AHC patient mutations disrupt cellular function beyond haploinsufficiency. Here, all eight heterozygous AHC model primary and replicate strain animals showed robust dominant NMJ defects in both aldicarb time-response and dose-response assays (Figs. 3, 4), as well as in levamisole time-response assays (except for E815K/+, Fig. 5). This is in stark contrast to heterozygous *eat-6(lf)/+* animals, which always showed normal sensitivity to aldicarb and levamisole (Figs. 3B,C, 4B,C, 5B,C). Given that heterozygous *eat-6(lf)/+* animals did not exhibit dominant NMJ defects, but all heterozygous AHC model animals did, we conclude that NMJ function in heterozygous AHC model animals must be disrupted through a mechanism beyond simple loss of pump function. Thus, we rule out haploinsufficiency as an explanation for NMJ defects in heterozygous AHC model animals.

Because heterozygous AHC model *C. elegans* have dominant NMJ defects that are not solely due to decreased pump function, the AHC mutations are classified as genetic “gain-of-function” alleles. By definition, genetic “gain-of-function” alleles can be hypermorphic (increased EAT-6 activity), antimorphic (dominant-negative, antagonising normal EAT-6 function), and/or neomorphic (novel activity, unrelated to normal EAT-6 function). Because homozygous *eat-6* decreased function alleles are hypersensitive to aldicarb (Doi and Iwasaki 2008; Sorkaç et al. 2016; Govorunova et al. 2010), a hypermorphic allele would be expected to cause aldicarb resistance. But, heterozygous AHC model animals are hypersensitive to aldicarb (Figs. 3,4). Thus, increased EAT-6 activity is an unlikely explanation for dominant NMJ defects in heterozygous AHC model animals. Results presented here cannot distinguish between “gain-of-function” antimorphic and/or neomorphic actions caused by AHC variants; either of these could potentially explain the observed dominant NMJ defects that go beyond haploinsufficiency or reduced EAT-6 function.

We see further evidence that AHC patient mutations can cause defects beyond loss of pump activity when considering the developmental defects of homozygous AHC model animals. Homozygous *eat-6(lf1)* animals have severe development defects that are similar to those observed in homozygous E815K animals. However, development defects in homozygous D801N and G947R animals were more severe than those in homozygous *eat-6(lf1)* animals (Fig. 2C,D). For D801N and G947R animals, this is further evidence that a mechanism beyond simple loss of ATPase activity is likely contributing to homozygous development defects.

### *C. elegans* AHC models have allele-specific impairments

All AHC alleles likely share a common disease mechanism, but there may be additional allele-specific defects. In AHC patients, there is a wide spectrum of symptom presentation and severity (Panagiotakaki et al. 2015; Viollet et al. 2015; Yang et al. 2014; Vezyroglou et al. 2022); the *C. elegans* AHC models also differ in their defects. Most notably, L839P is a comparatively weaker allele in patients (Pratt et al. 2020; Yang et al. 2014), and homozygous L839P *C. elegans* AHC model animals had the least severe homozygous survival and development defects (Fig. 2B-D). Despite exhibiting no growth past the L1 stage, homozygous L839P animals survived for at least 10 days while the vast majority of homozygous D801N, E815K, and G947R larvae died within 24 hours. For reference, under these conditions wild-type *C. elegans* reach the mid-L1 stage by 20 hours and reach adulthood within 3. Because the eight primary and replicate heterozygous AHC model strains were not examined in the same trials on the same days, it is difficult to directly compare the severity of dominant aldicarb response defects between the AHC models. However, dominant NMJ defects in primary and replicate L839P/+ model animals may be weaker (based on aldicarb time-response, Fig. 4K,L,Q).

While at least one strain from all four AHC model animals showed a dominant hypersensitivity to levamisole indicating post-synaptic body-wall muscle defects, the defect in heterozygous D801N/+ and maybe G947R/+ animals were arguably more severe. A levamisole response defect could only be detected in a paired t-test of median immobilization time for D801N/+, D801N*/+, and G947R*/+ model animals (Fig. 5). In humans, cardiac muscle impairment is observed much more frequently in patients carrying the D801N allele; D801N patients often have short QT syndrome and have an increased risk of life-threatening arrhythmias and sudden cardiac death (Moya-Mendez et al. 2021)*. C. elegans* do not have a heart, but their body-wall muscle shares key contractile properties with vertebrate cardiac muscle. It is interesting that the AHC patient mutation that most frequently causes cardiac muscle defects in patients is also the *C. elegans* model with the strongest and most reproducible post-synaptic muscle defect.

Allele-specific defects in the *C. elegans* AHC models are also apparent when sleep and arousal are examined. All *C. elegans* AHC model animals have decreased arousal thresholds during lethargus sleep bouts which suggests poor sleep quality, altered sensory circuit function, or an imbalance of neuromodulators (Cho and Sternberg 2014; Scheer and Bargmann 2023). However, only heterozygous G947R/+ animals showed an overall decrease in sleep quantity (Fig. 6). Additionally, only heterozygous E815K/+ model animals had a shorter L4/adult lethargus duration, decreased arousal thresholds in L4, and decreased wake bouts during lethargus which suggests difficulties transitioning between wake and sleep states or may indicate broader developmental defects.

Many factors could explain how different alleles in the same protein can lead to varying phenotypes and severities. The physical location of each point mutation within the Na^+^/K^+^ ATPase may affect the extent to which pump function is decreased. For example, D801N and G947R mutations are located in transmembrane domains near ion binding sites, while E815K and L839P mutations are located further away in intracellular loops (Li et al. 2015). Furthermore, alleles may induce different antagonistic pathways that depend on the extent of ion gradient disruption, cascading effects in the ER/Golgi (Arystarkhova et al. 2019, 2021; Arystarkhova and Sweadner 2024), or the role of the Na^+^/K^+^ ATPase in localization of post-synaptic receptors (Doi and Iwasaki 2008). We must consider the possibility that each AHC patient allele may respond differently to genetic modifiers or chemical intervention due to allele-specific impairments.

### Loss of *ncx-4* function inconsistently modified NMJ defects across alleles in *C. elegans* AHC models

The conservation of neurological genes and pathways as well as the ease of genetic manipulation in *C. elegans* mean that these AHC models may be a powerful tool for investigating AHC pathophysiology. Foundational work by the Palladino Lab in 2014 identified 33 genetic modifiers of a *Drosophila* G755S model of AHC. We found cross-species modification of AHC-related phenotypes for some AHC alleles when the function of a conserved NCKX was lost. *ncx-4* encodes a K^+^-dependent Ca^2+^/Na^+^ exchanger that is primarily expressed in adult neurons (Winkfein et al. 2004). Two different loss-of-function alleles in *ncx-4* suppressed dominant aldicarb dose-response defects in heterozygous G947R/+ animals However, only one of the two *ncx-4* loss-of-function alleles suppressed NMJ defects in heterozygous E815K/+ and L839P/+ animals. In all cases, the extent of suppression by *ncx-4* loss-of-function alleles was small. It is possible that *ncx-4* is a weak suppressor for all three AHC alleles, making it difficult to reproducibly detect subtle changes in aldicarb sensitivity. Or, it is possible that loss of *ncx-4* function only reliably modifies defects in G947R/+ animals due to allele-specific impairments.

In the *Drosophila* AHC model, heterozygous loss of *Nckx30C* function enhanced bang-sensitive paralysis defects, while homozygous loss of *ncx-4* function sometimes suppressed NMJ defects in the *C. elegans* AHC models. These are very different behaviors; *Drosophila* bang-sensitive paralysis behavior is regulated through different circuitry and pathways than *C. elegans* NMJ function. When assessing different behaviors in multiple species, the perturbation of a conserved gene could exacerbate one behavior and ameliorate another. Overall, if *ncx-4* is a cross-species modifier, we conclude that calcium homeostasis could play a role in AHC pathophysiology and that *C. elegans* AHC models can be used to identify conserved genetic mechanisms relevant to AHC. We plan to implement this same approach to test the remaining candidate modifier genres from the *Drosophila* screen in the *C. elegans* AHC models.

## Materials and Methods

### *C. elegans* strains and maintenance

All strains were reared at 20°C on standard Nematode Growth Medium (NGM) seeded with OP50 *E. coli*, unless otherwise indicated. All assays used age-matched hermaphrodites. See supplemental materials for the full strain list. Heterozygous AHC model and control strains were balanced by either the *tmC12[myo-2p::mCherry]* (Dejima et al. 2018) or *nT1[qIs51 (myo-2p::GFP, pes-10p::GFP, F22B7.9p::GFP)] (Ferguson and Horvitz 1985; Clark et al. 1988; Edgley et al. 2006)* balancer chromosomes. The *nT1* balancer became unreliable in progeny produced by older hermaphrodites and many balanced progeny lacked GFP expression. Thus, *tmC12* balanced strains were used unless otherwise stated in the figure legend. Heterozygous AHC model animals balanced over *nT1* or *tmC12* grew from egg to L4 roughly 12 hours slower than CRISPR control animals.

### Statistical analysis

All experiments were performed blinded to genotype and, when possible, treatment. Quantitative data was analyzed using GraphPad Prism 10. In figures, P values <0.05 are displayed and ns indicates P>0.05. Unrounded P values can be found in the Supplementary Materials. All independent trials are biological replicates conducted on different days, using different animals. Even when data is aggregated in figure panels for visual purposes, statistical comparisons are only made between strains tested in parallel on the same day.

Usually, balanced AHC model, CRISPR control, wild-type, and *eat-6(lf)* animals were tested together in aldicarb and levamisole assays. When figure panels were assembled, we separated results for loss-of function alleles, CRISPR controls, and AHC alleles. Therefore, sometimes the same balanced wild-type and CRISPR control animals are shown in multiple panels. Each trial has a unique name in the Supplementary Materials. For aldicarb and levamisole assays, we use two-tailed paired t-tests to control for day-to-day variation in culture conditions, environmental factors, and assay plate preparation.

### CRISPR-Cas9 microinjection to generate AHC model and CRISPR control strains

Basic Local Alignment Search Tool (BLAST) was used for amino acid alignment of human ATP1A3 (NP_689509.1) and *C. elegans eat-6* (B0365.3) in Fig. S1. AHC model (D801N, E815K, L839P, G947R) and control (D801D, E815E, L839L, G947G) strains were generated using CRISPR-Cas9 homology directed repair of endogenous *eat-6* with a single-stranded oligonucleotide template using previously published methods (Dokshin et al. 2018; Ghanta et al. 2021; Kim et al. 2014). Two Alt-R CRISPR-Cas9 sgRNA (Integrated DNA Technologies) were used to target either side of each edited region in *eat-6*. Repair templates were Alt-R HDR Donor Oligo ssODNs (Integrated DNA Technologies), containing the AHC missense mutation, silent edits to generate restriction sites, silent edits to destroy PAM sites, and ≥30 bases of unedited, flanking homology arms (Fig. S2 and Supplementary Materials). Editing was confirmed using both PCR genotyping and Sanger sequencing (Eurofins Genomics). Primary and replicate AHC model and CRISPR control strains (derived from different F1 animals post-CRISPR editing) were arbitrarily selected then backcrossed to wild-type animals at least twice before analysis.

### PCR genotyping

All primers were ordered from Integrated DNA Technologies. One*Taq* DNA Polymerase were used in all PCRs. The Supplementary Materials contain a full list of primers used to genotype AHC model and CRISPR control strains.

### Homozygous development and growth assessment

Ten to twenty gravid adults were picked to three freshly seeded NGM plates and allowed to lay eggs at 20°C for three hours; the total amount of eggs per genotype was counted. After 18 hours at 20°C, homozygous animals were categorized as: unhatched, dead L1, live L1, motile L1, or motile L1 expressing *lin-4p::GFP*. Live L1 animals had spontaneous nose or tail twitches or minimally twitched in response to gentle prodding with a platinum wire. Motile L1 animals showed muscle movement throughout their entire body and were able to crawl forward any distance either spontaneously or after gentle prodding. Dead L1 animals were hatched but did not respond to prodding. *myo-2p::mCherry* fluorescence from the *tmC12* balancer “+” could not be detected in the egg, so any unhatched eggs after 18 hours were assumed to be homozygous unbalanced animals. For the development index, each of the five categories was given a score of 0-5 from most to least severe. The number of homozygous animals in each category multiplied by the category score were summed and divided by the total number of homozygotes to produce a development index score for each trial. For each genotype, between 117 and 211 homozygous animals were scored across three biological replicates. Data was analyzed using a one-way ANOVA with Tukey’s multiple comparisons test between all possible pairs. Eighteen hours after an egg-laying, some homozygous L1 animals were immobilized with 2,3-Butanedione monoxime (BDM) and mounted on 2% agarose pads. Images were captured under the 40x objective on a Zeiss AxioImager ApoTome and AxioVision software v4.8.

### Pharyngeal Pumping

Animals were picked at the L4 stage and assayed as adults one or eight days later. Videos of individual animals were recorded for 10 seconds on a PointGrey camera using FlyCapture 2. Full or partial grinder movement in any direction was manually counted as a pharyngeal pump. The number of pumps per 10 second video was counted and calculated into pumps/minute. Between 26 and 30 total animals were recorded across three biological replicates. Data was analyzed using a one-way ANOVA with Šídák’s multiple comparisons test between each AHC model and its respective CRISPR control strain.

### Egg laying

Eight age-matched day 1 adults laid eggs on freshly seeded NGM plates for six hours at 20°C and the total number of eggs were counted. Data was collected from three biological replicates and analyzed using one-way ANOVAs between +/+, two replicate AHC model strains, and their two replicate CRISPR control strains. A Šídák’s multiple comparisons test was used between each +/+ and CRISPR control strains, and between each AHC mutant and its respective CRISPR control. Additionally, paired t-tests were performed between N2 and *eat-6(ad467)* animals and between *+/+* and *eat-6(lf2)/+* animals.

### Growth on DA837 and HB101 *E. coli*

Bacterial lawns of DA837, HB101, or OP50 *E. coli* were seeded onto NGM plates and left to dry overnight before introduction of age-synchronized L1 animals from a bleach prep (Porta-de-la-Riva et al. 2012). The growth rates, locomotion, and overt behavior of heterozygous and homozygous AHC model animals were qualitatively compared.

### Long-term temperature stress

Animals were reared at 12°C and 27°C for at least three generations and observed for overt locomotion defects or changes to survival, growth, and development.

### Acute Heat-shock

Twenty animals per genotype were heat-shocked at 35°C for 2.5 hours on seeded NGM plates placed agar-side up on a heat block and then moved to 20°C for recovery. 57 to 64 hours later, survival was scored based on pharyngeal pumping and spontaneous or evoked motion. Data was collected from three biological replicates with 18-20 animals per genotype per trial. One-Way ANOVA with Šídák’s multiple comparisons test between CRISPR controls and AHC model animals.

### Acute Cold-shock

Twenty animals per genotype were cold-shocked at either 4°C for 17 hours or 0°C for 4 hours. At 4°C, seeded NGM plates were placed in the refrigerator in a singular layer. At 0°C, plates were individually sealed in parafilm and buried in ice in a single layer. After the cold-shock, plates were incubated at 20°C for 18 hours. After 18 hours, survival was scored based on pharyngeal pumping and spontaneous or evoked locomotion. Data was collected from three biological replicates, each with 20 animals per genotype.

### Aldicarb dose-response

The aldicarb dose-response assay was adapted from previous work (Nonet et al. 1997). One day prior to the assay, L4 hermaphrodites were selected and allowed to grow overnight on normal seeded NGM plates. NGM plates containing 0, 0.25, 0.5, 0.75, and 1 mM of aldicarb were prepared and left to dry on the benchtop overnight. On the day of the assay, a copper ring was melted onto the center of each aldicarb plate, and 10 µL of OP50 was seeded in the center of the ring (Hawkins et al. 2015). Once the OP50 dried, 10-15 day 1 adult animals of each genotype were transferred onto aldicarb plates of each dose. The plates were left for five hours at room temperature on the benchtop. After five hours, immobilization was scored by gently tapping animals twice on the head and twice on the tail with a platinum wire. If animals did not move, twitch, or pump after three seconds of observation, they were scored as immobilized. Escaping animals were censored. After five biological replicates, immobilization curves were analyzed using two-way ANOVA. Area under the curve values were calculated and then analyzed in paired t-tests. Only strains tested in parallel on the same days were compared.

### Aldicarb time-response

The aldicarb time-response assay was adapted from previous work and largely undertaken as described above for the dose-response assay. However, only 1 mM aldicarb (Mahoney et al. 2006) NGM plates were used and immobilization was assessed every hour for six hours. Four biological replicates with 10 to 22 animals per genotype per replicate were analyzed using a Mantel-Cox log-rank test for two strains tested in parallel. The hour when at least 50% of animals were first immobilized was recorded and analyzed in paired t-tests for two strains assayed in parallel. If a strain did not reach 50% immobilization by the final six hour timepoint, the 50% immobilization time was arbitrarily set as seven hours. Only strains tested in parallel on the same days were compared.

### Levamisole time-response

The levamisole time-response assay was adapted from previously described methods (Gottschalk et al. 2005; Hawkins et al. 2015; Doi and Iwasaki 2008) to be conducted similarly to the aldicarb time-response assay described above. Immobilization was examined on 100 uM levamisole NGM plates every hour for six hours. Three biological replicates with 9 to 27 animals per genotype per replicate were analyzed using a Mantel-Cox log-rank test performed between two strains tested in parallel on the same days.

### Developmentally timed sleep

Animals were visually staged at L4.2 (Mok et al. 2015) and imaged for twelve hours, including L4/A lethargus in individual polydimethylsiloxane (microfluidic chips containing 10 chambers, as described previously (Huang et al. 2017)) chambers. Images were taken every 10 seconds, and image subtraction was used to detect lack of motion as previously described (Huang et al. 2017). Up to four genotypes were tested per imaging session, and controls were pooled with other strains run within the same twenty-four-hour period. Each genotype was run in at least three sessions. In total, 22 to 34 animals were examined per genotype. Comparisons between AHC model animals and appropriate CRISPR controls used paired t-tests.

### Arousal

With modifications, arousal threshold assays were performed as previously described (Huang et al. 2017). Five to ten gravid adult animals were transferred to a freshly seeded NGM plate and placed at 20°C for forty-eight hours. Plates were then left under a dissection microscope light for thirty minutes to acclimate animals to the temperature and light. Plates were viewed at 4x magnification using a dissection microscope with the lid on. A 5 mW, a 405 nm laser affixed with a ∼0.5 mm pinhole cover was mounted to a magnetic manipulandum arm and positioned to emit light to the center of the microscope field of view. The tip of the laser pointer was fixed 3 cm above the lid of the plate as vertically as the scope permitted. Current was maintained at 0.28 amperes to ensure a consistent laser intensity of approximately 100 lux. Sleeping and awake animals were positioned with their pharynx in the center of the field of view, and the laser was activated until animals responded by moving one body bend forwards or backwards. In moving animals, a change in direction or an acceleration by one body bend was considered a response. Laser illumination was terminated when a response was detected, and response latency times and the direction of response (forward or reverse) were digitally recorded. Animals were visually staged as L4.2 (L4), L4.6-L4.8 (sleep bout, L4/A), L4.6-L4.8 (wake bout, L4/A), and vulva-everted young-adults (adult). Sleep was defined as absence of moving and feeding behavior. In total, thirty animals of each genotype at each stage were assayed in three biological replicates. Data was analyzed using a one-way ANOVA with Sidak’s multiple comparisons test between +/+ and *goa-1(lf)*, +/+ and CRISPR control, and CRISPR control and AHC model strain. A separate ANOVA test was used for each AHC model strain and its corresponding controls that were assayed together.

## Supporting information

Strains, Oligos, Raw Data

## Acknowledgements

We gratefully acknowledge support from Hope for Annabel, Hope4Livi, Cure AHC, and the AHC Foundation. Advice from Dr. Natalia Morsci, Nina Frost, Dr. Kathy Sweadner and other AHC researchers was invaluable. Some strains were provided by the CGC, which is funded by NIH Office of Research Infrastructure Programs (P40 OD010440). Some strains were provided by the National BioResource Project and National BioResource Project (NBRP) for the Experimental Animal “Nematode C. elegans” and are subject to a materials transfer agreement.

## Data and Resource Availability

All relevant data and details of resources can be found within the article and its supplementary information.

## Supplementary Figures PDF

**Supplemental Figure 1:**
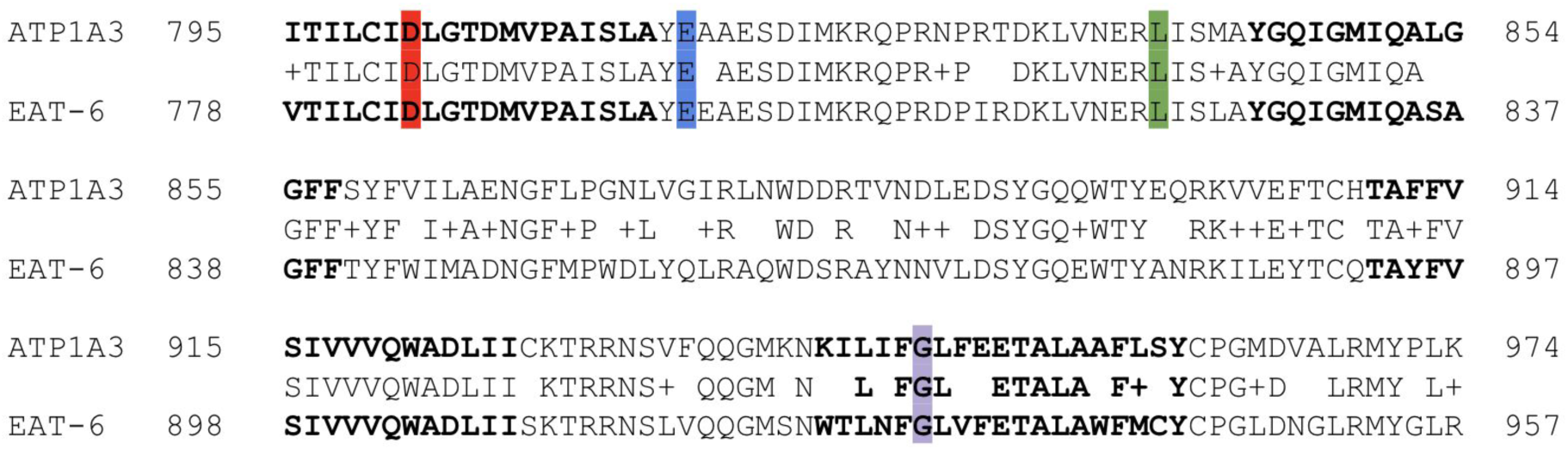
Partial alignment of Human ATP1A3 and C. elegans EAT-6. Conserved amino acid residues shown between the two protein sequences for human ATP1A3 (NP_689509.1) and *C. elegans EAT-6* (B0365.3); similar amino acids indicated with “+”. Bold indicates transmembrane domains 6 through 9. Highlighted residues indicate location of patient missense mutations inserted into endogenous *eat-6* via CRISPR/Cas9 to create *C. elegans* AHC models for D801N (red), E815K (blue), L839P (green), or G947R (purple).

**Supplemental Figure 2:**
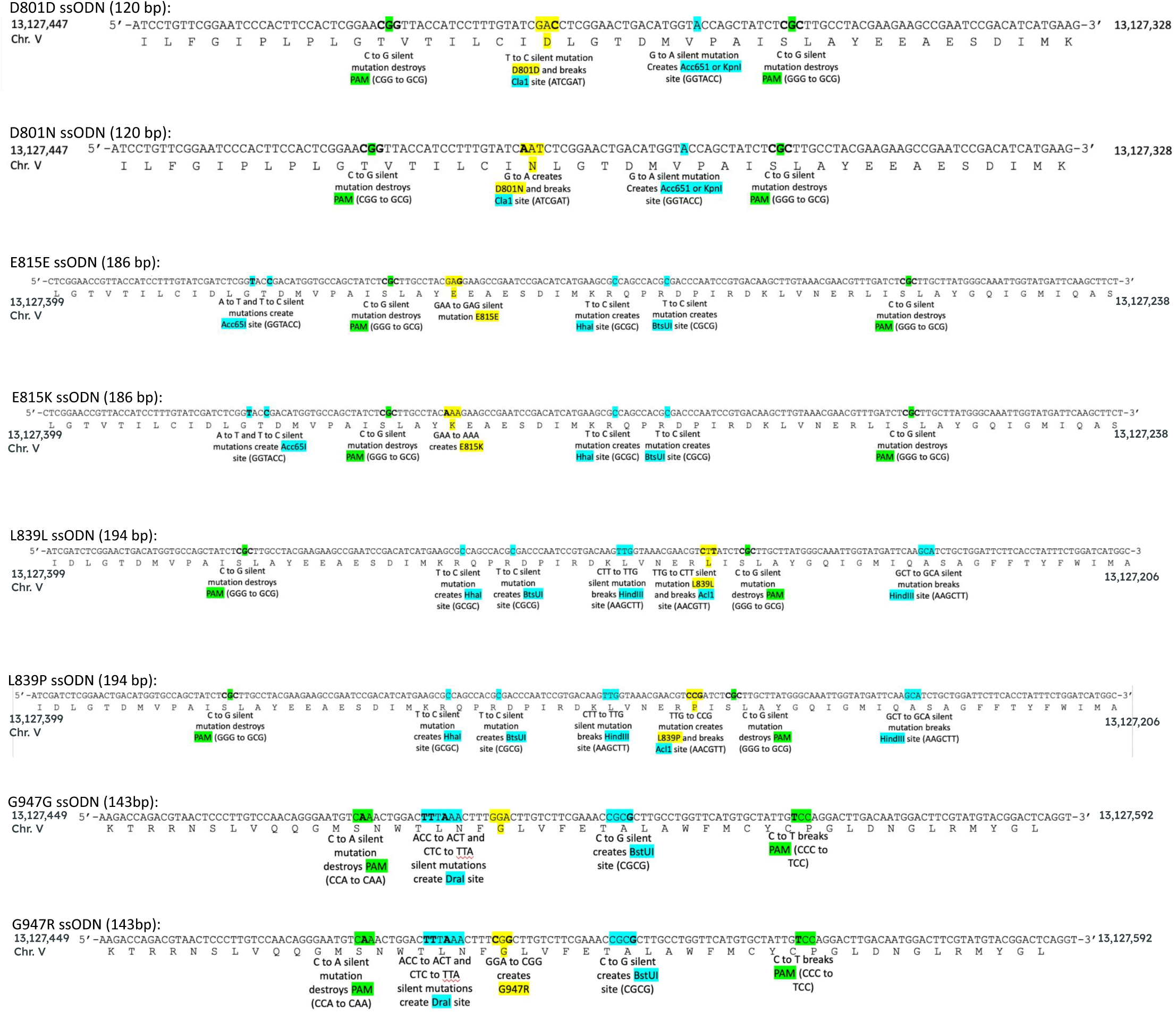
ssODNs used for generation of each AHC model and CRISPR control strain. DNA sequence of each ssODN is displayed above its translated amino acid sequence. Location on chromosome is noted. Each ssODN contains 30bp of unedited flanking homology arms, silent mutations to destroy sgRNA PAM sites (green), AHC patient missense mutation for AHC model strains and CRISPR control silent mutations for control strains (yellow), and silent mutations to introduce or remove restriction sites for PCR-based genotyping (blue).

**Supplemental Figure 3:**
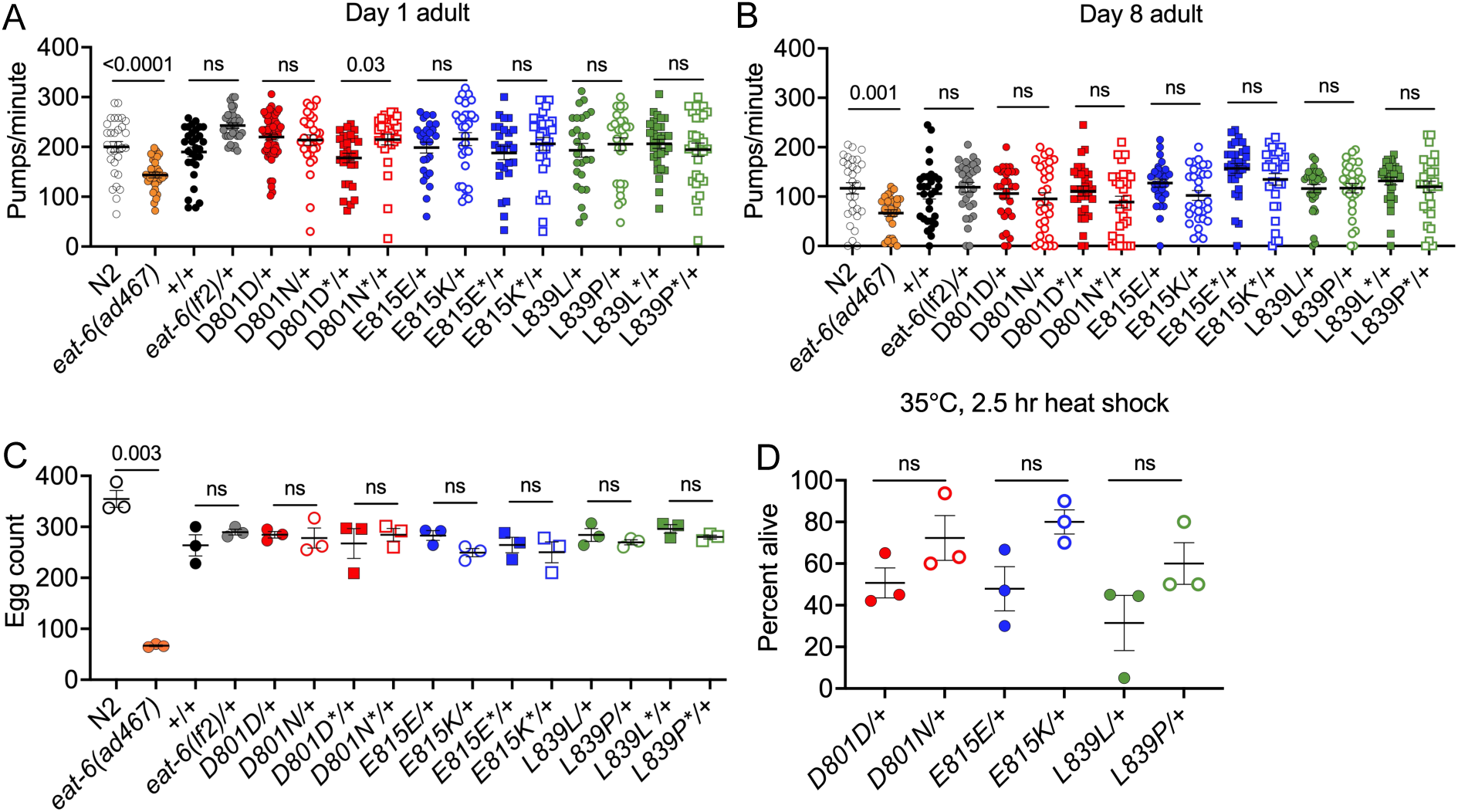
Pharyngeal pumping, egg-laying, and stress-response of AHC model animals. Pharyngeal pumps per minute in day 1 adult (A) and day 8 adult (B) animals. Data accumulated from three trials, n=26-30 animals per genotype. N2, *eat-6(ad467), +/+, eat-6(lf2)/+* tested in a one-way ANOVA with Šídák’s multiple comparisons test between shown pairs. One-way ANOVA of +/+, the heterozygous primary and replicate CRISPR control strains, and heterozygous primary and replicate AHC model strains tested in with Šídák’s multiple comparisons test between each CRISPR control and AHC mutant pair (shown on graph) and +/+ and each CRISPR control (not shown on graph P>0.05). For day 1 adults, +/+, D801D/+, D801D*/+, D801N/+, D801N*/+ were assayed separately from the other genotypes, but data is combined for presentation in A. Statistics were performed only on animals tested in parallel. (C) Number of eggs laid by 8 adults in 6 hours. Three biological replicates. One-way ANOVAs between +/+, two replicate AHC model strains, and their two replicate CRISPR control strains with Šídák’s multiple comparisons test between +/+ and each CRISPR control strain, and between each AHC mutant and its respective CRISPR control. This analysis was performed for each AHC allele set. Comparisons between +/+ and CRISPR control strains not shown on graph (all P>0.05 except +/+ vs E815E*/+ P=0.0020). Unpaired t-tests with Welch’s correction between N2 and *eat-6(ad467)* and between *+/+* and *eat-6(lf2)/+*. (D) Percent of heterozygous adult animals alive 60 hours after a 2.5 hour, 35°C heat-shock. n=18-20 animals per genotype per trial. Three biological replicates. One-Way ANOVA with Šídák’s multiple comparisons test between shown pairs. In all panels, One “+” in heterozygous animals represents the *nT1* balancer, error bars indicate mean +/- SEM, ns P>0.05.

**Supplemental Figure 4:**
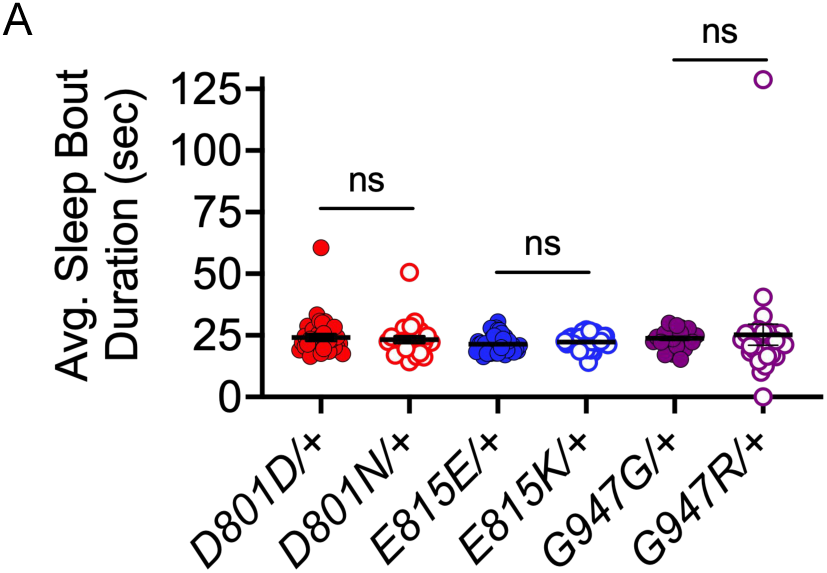
Average sleep bout duration in heterozygous AHC model animals. Average sleep bout duration for D801N/+, E815K/+, and G947R/+ model animals and corresponding CRISPR control strains using Welch’s unpaired t-test. n= 34-21 animals per genotype. “+” chromosome in D801D/+, D801N/+, E815E/+, and E815K/+ animals is the *nT1* balancer. “+” chromosome in G947G/+ and G947R/+ animals is the *tmC12* balancer. Error bars represent mean +/- SEM.

**Supplemental Figure 5:**
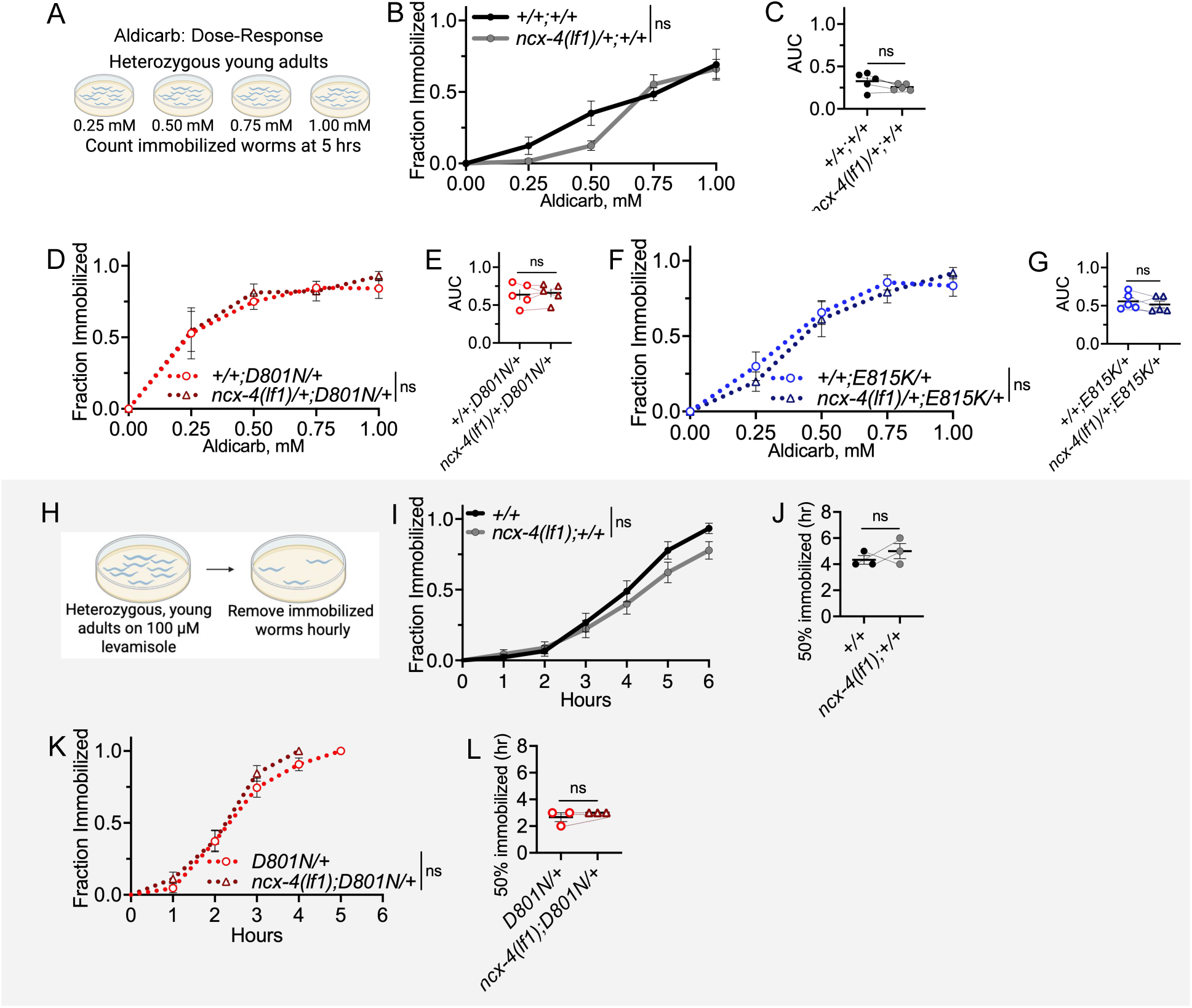
Heterozygous loss of ncx-4 did not alter AHC model aldicarb defects, and homozygous loss of ncx-4 did not suppress D801N/+ levamisole defects. (A) Aldicarb dose-response assay. The number of young-adult animals immobilized on 0, 0.25, 0.5, 0.75, and 1 mM aldicarb was recorded at five hours. (B) Dose-response curve for *ncx-4(tm5106)/+;+/+* and +/+ animals. One “+” chromosome is the *tmC18* balancer, and one “+” chromosome is the *tmC12* balancer in all strains used in B-G. Two-way ANOVA. (C) Area under the curve (AUC) from each trial in B, Paired t-test. (D) Dose response curves for D801N/+ animals with and without one copy of *ncx-4(tm5106).* Two-way ANOVA. (E) AUC from each trial in D. Paired t-test. (F) Dose response curves for E815K/+ animals with and without one copy of *ncx-4(tm5106).* Two-way ANOVA. (g) AUC from each trial in F. Paired t-test. For all two-way ANOVA analysis, the displayed P value results from genotype as the source of variation. Data for each genotype results from five biological replicates. For each trial, n=13-15 animals per genotype. (H) Levamisole time-response assay. The number of heterozygous, young-adult animals immobilized on 100 uM levamisole was recorded hourly for six hours. (I) Immobilization curve for *ncx-4(tm5106);+/+* and +/+ animals. In all strains in I-L, one “+” chromosome is the *tmC12* balancer. Mantel-Cox log-rank analysis. (J) Median immobilization time for each trial in I. Paired t-test. (K) Immobilization curve for D801N/+ animals with and without one copy of *ncx-4(tm5106).* Mantel-Cox log-rank test. (L) Median immobilization time for each trial in K. Paired t-test. Data for each genotype is from three biological replicates. For each trial, n=13-15 animals per genotype. All error bars indicate mean +/- SEM, ns indicates P>0.05.

**Supplemental Figure 6:**
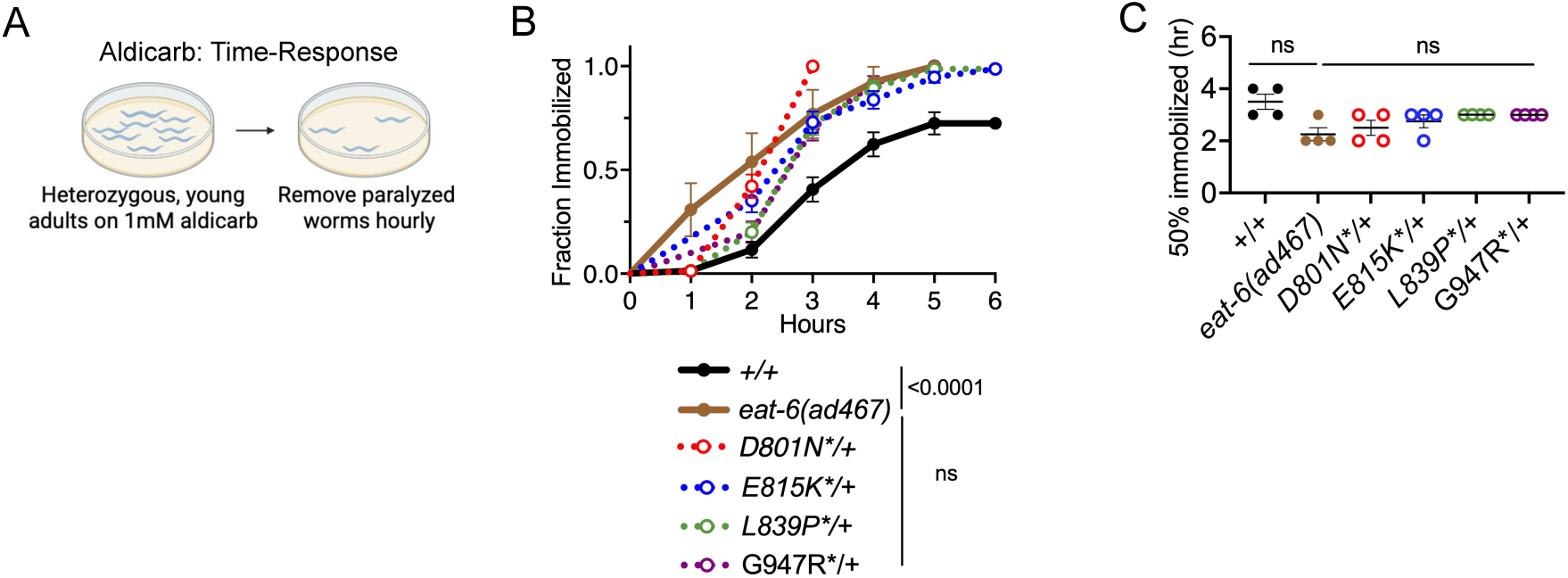
Dominant NMJ defects in AHC model animals are similar in magnitude to eat-6(ad467) animals: aldicarb time-response. (A) Aldicarb time-response assay. The number of heterozygous, young-adult animals immobilized on 1 mM aldicarb was recorded hourly. (B) Immobilization curves for wild-type +/+, homozygous *eat-6(ad467)* partial loss-of-function, and heterozygous AHC model D801N/+, E815K/+, and L839P/+ animals. Mantel-Cox log-rank analysis performed between +/+ and *eat-6(ad467)* animals, and between *eat-6(ad467)* and each AHC model strain. (C) Median immobilization time for each trial in B. Paired t-test between between +/+ and *eat-6(ad467)* animals, and between *eat-6(ad467)* and each AHC model strain. Each genotype tested in four biological replicates. For each trial, n=10-22 animals per genotype. Error bars indicate mean +/- SEM. ns indicates P>0.05. In all strains except *eat-6(ad467),* one “+” represents the *tmC12* balancer. Data for +/+, D801N*/+, E815K*/+, L839P*/+, and G947R*/+ is re-used from Figure 4. Note that G947R*/+ model animals animals were assessed in separate trials but are presented together for visual purposes to compare relative magnitude of aldicarb hypersensitivity.

